# Distinctive mechanism of LHCSR3 expression and function under osmotic stress in *Chlamydomonas reinhardtii*

**DOI:** 10.1101/2023.05.12.540499

**Authors:** Sai Kiran Madireddi, Ranay Mohan Yadav, Pushan Bag, Mohammad Yusuf Zamal, Rajagopal Subramanyam

## Abstract

Light-harvesting complex stress-related protein 3 (LHCSR3) expression is observed in various protoxidizing conditions like high light and nutrient starvation. LHCSR3 expression is essential for energy-dependent quenching (qE), whereas its role under nutrient starvation is elusive. It is also unclear how nutrient starvation can induce LHCSR3 expression under subsaturating light intensities. To study the role of LHCSR3 under nutrient starvation, the *C. reinhardtii* cells are grown under osmotic stress that would prevent water uptake; therefore same holds true for soluble nutrients in the medium. In this work, we have shown that LHCSR3 expression can occur under osmotic stress and subsaturating light intensities, whereas it does not elicit qE. Further examination of thylakoid membrane architecture from wild-type and *npq4* mutant grown under nutrient starvation revealed that LHCSR3 expression affects the interaction between the PSII core with its peripheral LHCII antenna and possibly can prevent excitation energy transfer. Thylakoid lumen acidification is essential for the expression and function of LHCSR3. Under saturating light intensities, this is achieved by the increased rate of photosynthetic electron flow coupled with proton translocation into the thylakoid lumen. Whereas, under nutrient starvation, the reports of LHCSR3 expression also showed reduced photosynthetic electron flow. Therefore, an alternative mechanism should exist for developing the proton gradient. We observed the downregulation of chloroplast (cp) ATP synthase activity and its abundance under osmotic stress, suggesting the role of (cp) ATP synthase in thylakoid lumen acidification under reduced photosynthetic electron flow. This observation is supported by the expression of LHCSR3 in (cp) ATP-synthase mutant *atpF* upon exposure to moderate light intensity. This study proposes that the mechanism of LHCSR3 expression and its functionality can vary with the type of photooxidizing stress.

## Introduction

Optimum photosynthesis requires a sensitive balance between photoreception and its utilization. This involves exposure to optimum light intensity and a constant supply of nutrients.

For photosynthesis to function at maximum efficiency, it requires a constant supply of nutrients like carbon, nitrogen, oxygen, and minerals that contain certain metal ions that constitute co-factors for photosynthetic reactions. The limited supply of these nutrients can lead to compromised photosynthetic activity (Yruela, 2013) due to the alteration of the balance between the absorption of light energy and carbon fixation reactions. This imbalance in photosynthetic activity can result in ROS generation that can prevent the repair of PSII, causing photoinhibition (Murata et al., 2007). Most of the minerals are absorbed by the plants in water-soluble form. Consequently, any change in water availability affects the plant’s capability to receive nutrition. Light and nutrient supply fluctuate in natural conditions, which are deleterious to photosynthetic machinery. These environmental fluctuations can lead to an imbalance between photoreception and its utilization. Therefore, photosynthetic organisms must adapt and/or acclimate to environmental changes to sustain photosynthetic activity and prevent photoinhibition (Murata et al., 2007). All photosynthetic organisms have evolved with mechanisms to dissipate excess energy absorbed by photosynthetic machinery, collectively called non-photochemical quenching (NPQ). NPQ has different components based on the induction and relaxation kinetics; fast and reversible quenching (qE) (Muller et al., 2001), state transition quenching (qT) (Minagawa, 2011), Zeaxanthin dependent quenching (qZ) (Nilkens et al., 2010), photoinhibitory quenching (qI) (Krause, 1988) and recently termed an alternative form of quenching called sustained quenching (qH) (Malnoë, 2018). qE is the fastest and most studied component of NPQ. Under high light, qE is induced within a tenth of a second and can relax within minutes on exposure to dark or sub-saturating light intensity (Niyogi, 2000). In green algae, *C. reinhardtii*, the induction of qE requires gene products of light-harvesting complex stress-related (LHCSR) *LHCSR1, LHCSR3.1/3.2* and *PSBS* (Peers et al., 2009; Correa-Galvis et al., 2016). LHCSR isoforms bind to Chl *a*, Chl *b*, lutein and violaxanthin/zeaxanthin, in addition to proton binding residues on the luminal side (Bonente et al., 2011). The sensitivity of qE to uncopular nigericin (Tokutsu and Minagawa, 2013) emphasizes the importance of ΔpH for PSBS and LHCSR3 to function (Tian et al., 2019).

Research on LHCSR3 in the past decade has provided substantial information on its expression, physical association with photosystems, and qE induction properties under high light (Peers et al., 2009); blue light (Petroutsos et al., 2016) and UV light (Allorent et al., 2016). In addition to high-light induced expression, several genome and proteome-based approaches have reported the expression of LHCSR3 in *C. reinhardtii* cells grown under Iron, Phosphorous, Copper and Sulphur starvation (Im et al., 2003; Zhang *et al*., 2004; Moseley et al., 2006; Naumann et al., 2007; Strenkert et al., 2016). It is noteworthy that LHCSR3 expression during nutrient starvation occurs under moderate light intensity, and it is not clear regarding the role of LHCSR3 under nutrient starvation. To clarify the role of LHCSR3 under nutrient starvation, here, in this work, we employed Polyethylene glycol-8000 (PEG-8000) as an osmoticum that can limit water uptake by Chlamydomonas cells that can result in nutrient starvation. PEG is non-phytotoxic and is known to lower the osmotic potential in nutrient solutions (Lawlor, 1970). We considered this growth condition as nutrient starvation due to osmotic stress mimicking drought stress in higher plants in terms of nutrient uptake. This study shows that the LHCSR3 expression occurs in the *C. reinhardtii* cells grown under osmotic stress with moderate light intensity and it does not result in qE upon high light exposure. Using lpBN-PAGE and circular dichroism studies it is observed that LHCSR3 expression is correlated with the destabilization of PSII-LHCII complexes that could possibly hinder excitation energy transfer from the LHCII antenna to the PSII reaction center thus can avert photodamage.

Another question we tried to address in this work is regarding the possible retrograde signaling pathway for LHCSR3 expression under osmotic stress. Recent reports show the involvement of the pathways that consist of the photoperiod signaling system CONSTANS (Tokutsu et al., 2019), E3 ubiquitin ligase complex (Gabilly et al., 2019), calcium sensor protein (CAS) (Petroutsos *et al*., 2011), photoreceptors like UVR8 (Allorent et al., 2016) and flavin-containing phototrophins (PHOT) (Petroutsos et al., 2016). The involvement of the CAS pathway and the presence of Ca^2+^/H^+^ antiporter in the thylakoid membrane suggests the role of light-driven uptake of Ca^2+^ for the expression of LHCSR3 (Ettinger et al., 1999; Petroutsos et al., 2011) and this link is not established so far. If this holds true, we suppose that the LHCSR3 expression and function depend on thylakoid lumen acidification. As the rate of proton translocation into the thylakoid lumen is proportional to the rate of electron transport, and nutrient starvation negatively affects the photosynthetic electron transport rate (Zhang et al., 2004; Moseley et al., 2006; Terauchi et al., 2010), the ΔpH across the thylakoid membrane could be low. This condition raises the question about the possible ways to generate ΔpH across thylakoid membranes under low electron transport rate. With this study, we propose a model where the development of ΔpH across the thylakoid membrane under moderate light intensity and osmotic stress (nutrient starvation) can be attributed to the compromised function of the (cp) ATP synthase complex.

## Materials and Methods

### Culture and growth conditions

Chlamydomonas wt strain CC-125, *npq4* mutant (when indicated) was grown in TAP medium supplemented with different concentrations of PEG - 8000 (Sigma - Aldrich, Germany). Under continuous illumination (50 μmol photons m^-2^ s^-1^) using white fluorescent light with 5000 K spectrum, shaking at 120 rpm. Cells were grown until the control cell’s growth reached OD 1 at 750 nm. OD was measured in Perkin Elmer Lambda 25 UV/VIS spectrophotometer using 1 cm pathlength cuvette.

### Visualization of cell morphology

After growing under different concentrations of PEG, cells were spotted on a polylysine-coated slide for imaging. Cells are visualized using a TCS SP8 confocal laser-scanning microscope (Leica) at RT (25 °C). Imaging settings are as follows: images were taken by scanning mode using LASX software and a × 63 numerical aperture 1.4 oil objective, with 16-line averaging and 2-frame accumulation. For Chlorophyll, autofluorescence excitation/ emission settings are 561 nm/ 650 – 700 nm with HyD SMD hybrid detector.

### Growth stage and LHCSR3 expression measurements

Chlamydomonas wt strain CC-125 was grown in TAP medium until growth reached OD 1 at 750 nm. These cells were re-inoculated in a fresh TAP medium supplemented with 0% PEG (control) and 7.5% PEG (PEG) to the initial concentration of OD 0.2 at 750 nm. These cells were grown for 72h at 25 °C (± 2 °C) under continuous illumination (50 μmol photons m^-2^ s^-1^) using white fluorescent light with 5000K spectrum, shaking at 120 rpm, and samples were collected at every 6 h intervals to study OD at 750 nm, OD was measured with UV/VIS spectrophotometer (PerkinElmer Lambda 35) using 1 cm pathlength cuvette. The samples collected were used to study LHCSR3 expression using immunoblots.

### Chl fluorescence measurements

Chlorophyll fluorescence was measured using a pulse-amplitude-modulated fluorometer DUAL-PAM-100 (Walz) for RLCs and NPQ measurements and Handy PEA (Hansatech Instruments) for FLJ, F_m_ and F_v_/F_m_ determination. For light curve analysis, Chlamydomonas cells were dark-adapted for 15 min, and rapid light curves (RLC) were recorded with increasing light intensities. The light response curves recorded for ETR (II), Y (PSII), Y (NPQ), qP and Y (NO) were calculated and obtained from waltz software. For NPQ measurements, Wt *C. reinhardtii* cells (CC-125) were grown in TAP medium under growth light (∼50 μmol photons m^-2^ s^-1^), HL (∼500 μmol photons m^-2^ s^-1^) and osmotic stress with PEG supplemented TAP medium under growth light for 72 h. After dark adaptation, cells were pre-illuminated for 2 min with a weak (3 µmol photons m^−2^ s^−1^) far-red LED; maximum fluorescence (F_m_) and changes in maximal fluorescence in light (F_m_’) are measured by applying saturating pulse. NPQ was calculated using the formula NPQ = (F_m_-F_m_’)/ F_m_’; actinic light was 660 µmol photons m^−2^ s^−1^ and saturating light, 4,080 µmol photons m^−2^ s^−1^. The far-red LED was kept on during dark recovery to oxidize PSI and prevent over-reduction of the PQ pool (Bonente et al., 2011).

### Photoinhibition experiment

Wild type Chlamydomonas cells grown in TAP medium and TAP medium supplemented with PEG-8000 at 50 μmol photons m^-2^ s^-1^. These cells were incubated in the dark for 30 min in the presence of chloramphenicol (100 µg ml^−1^) and then exposed to strong fluorescent light (500 μmol photons m^-2^ s^-1^) for 1 h. The cells were then centrifuged and stored at −20 °C for further immunoblotting with an anti-D1 antibody.

### Isolation of thylakoids

Thylakoids were isolated as previously described (Kargul et al., 2003; Subramanyam et al., 2006) with minor modifications (Madireddi et al., 2019). Chlamydomonas cells were harvested from suspension cultures by centrifugation at 1000 x g for 5 min. The cell pellet is washed three times with PBS (phosphate buffer saline) for 3 min at 1000 x g to remove traces of PEG-8000 from the cells. The cell pellet was suspended in 25mM HEPES (pH 7.5, KOH), 0.3M sucrose, 10 mM MgCl_2_, 5 mM CaCl_2_, 10 mM NaF and 1 mM PMSF and homogenized by sonication. Homogenate was centrifuged at 1000×*g* for 3 min to remove unbroken cells and starch. The supernatant was collected in fresh tubes and centrifuged at 10000×*g* for 10 min. The pellet was suspended in 5mM HEPES (pH 7.5, KOH), 0.3M sucrose, 10mM NaF and 10 mM EDTA. This suspension was centrifuged at 18,000×*g* for 10 min. The resultant thylakoid pellet was suspended in 20mM Tricine (pH 7.5), 0.3M sorbitol, 10mM MgCl_2_, 10mM NaF, and 5mM CaCl_2_ and stored at −20 °C until further use. All the steps of isolation are carried out under 4 °C.

### SDS-PAGE and immunoblot analysis

The chlorophyll content of the cells was estimated using 80% acetone (Porra et al., 1989). Total cellular proteins were solubilized in the SDS-PAGE sample buffer (2% SDS and 0.1 M dithiothreitol) at 100 °C for 1 min. Samples were loaded with equal chlorophyll concentration (1 µg) per well, and proteins were separated in SDS/urea–PAGE (15% polyacrylamide, 6 M urea) (Laemmli, 1970) using mini-protean apparatus (Bio-Rad USA). These separated proteins were transferred to a nitrocellulose membrane using trans-blot apparatus (Bio-Rad USA). These blots were blocked with 4% skimmed milk in 1X TBST (for anti-phospho threonine antibody 4% BSA in 1X TBST) and probed with primary antibodies against photosynthetic proteins. Antibodies for photosynthetic proteins were diluted per manufacturer instructions (Agrisera Sweden), PsaA (1:3000). PsbD D2 protein of PSII (1:5000), PsbA D1 protein of PSII (1:10000), LHCSR3 (1:1000), AtpC (1:10000), Lhcb2 (1:5000), Lhcbm5 (1:5000), PetA (1:5000), STT7 (1:1000) and anti phosphothreonine (Cell Signalling Technology) (1:5000). Anti-rabbit-HRP linked antibody (1:10000) is used as the secondary antibody (Cell Signalling Technology). Chemiluminescence signals were detected using an ECL reagent (Thermo Fisher Scientific) by Chemidoc-touch (Bio-rad USA), and images were analyzed with image-lab software (Bio-rad USA).

### Circular Dichroism spectra measurements

Circular Dichroism (CD) spectra of *C. reinhardtii* cells were recorded by JASCO 810 spectropolarimeter. Measurements were carried out at 25 LJC, between 400 and 800 nm at 100 nm/min scan speed using 3 nm band-pass and 2 nm step size in a cuvette with a 1 cm optical path length with 3 spectral accumulations. Chlorophyll concentration is adjusted to 20 μg/mL as per Subramanyam et al., 2006.

### lpBN-PAGE

Photosynthetic membrane complexes were separated from intact thylakoid membranes by Large Pore Blue Native gel (lpBN) systems as described earlier (Järvi et al., 2011). Acrylamide gradient of 3.5–12.5% (w/v) T and 3% (w/v) C in the separation gel and 3% (w/v) T and 20% (w/v) C in the stacking gel was used. Samples for lpBN-PAGE were prepared (Schagger & Vonjagow, 1991) using 1% β-DM (β-D-maltoside) to solubilize the thylakoid membranes. This step is followed by separating complexes under constant current at 4 °C (Sirpiö et al., 2011). The second dimension of denaturing SDS-PAGE is employed to identify bands obtained in lpBN-PAGE. The strips from the first dimension were excised and solubilized in denaturing Laemmli buffer for one hour with gentle shaking [138 mM Tris/HCl (pH 6.8), 6 M urea, 22.2% (v/v) glycerol, 4.3% (w/v) SDS and 5% (v/v) 2-mercaptoethanol] for 1 h at 20°C. After solubilization, the strips were loaded on top of SDS/PAGE gel (15% (w/v) polyacrylamide and 6 M urea) and sealed with 0.5% agarose in SDS/PAGE running buffer. This is followed by electrophoretic separation of the protein subunits from the complexes at a constant voltage. After electrophoresis, the proteins were visualized by Coomassie Blue staining. Identification of photosynthetic supercomplexes in different bands obtained from lpBN-PAGE and second dimension PAGE was performed by immunoblotting with antibodies specific to PsbA (1:10000), PetA (1:5000), PsaA (1:3000) and LHCSR3 (1:1000). All antibodies were purchased from Agrisera (Sweden).

### ROS measurements

Intracellular ROS is measured by using an oxidant-sensing fluorescent probe 2,7-dichlorodihydrofluorescein diacetate (H2DCFDA) (Sigma-Aldrich) in the method described (Rastogi et al., 2010). Chlamydomonas cells grown under different conditions are collected by centrifugation, washed with a fresh TAP medium to remove PEG, and then exposed to light (50 μmol photons m^-2^ s^-1^) for 15 min. These cells are suspended in 1X PBS containing 5 µM H2DCFDA and incubated for 30 min in the dark with gentle shaking. After incubation, samples were washed three times with 1X PBS. Images were captured using Carl Zeiss NL0 710 Confocal microscope. H2DCFDA fluorescence was detected in a 500 – 530 nm bandpass optical filter with an excitation wavelength of 492 nm and an emission wavelength of 525 nm. Chlorophyll auto-fluorescence was detected using a long-pass filter of 600 nm. Samples were viewed with a 60× oil immersion lens objective by using the ZEN, 2010 software. ROS was quantified according to the method described by (Chouhan et al., 2022) using a spectrophotometer. At equal chlorophyll concentration, fluorescence was measured using a Microplate reader (Tecan M250) at an excitation of 485 nm and an emission of 530 nm. We calculated the total ROS by subtracting the value of fluorescence with dye and without dye.

### Electrochromic shift analysis

Rapid relaxation kinetics induced by a single turnover saturating flash was recorded at 515 nm from intact *C. reinhardtii* cells using DUAL PAM-100 (Walz, Effeltrich, Germany) equipped with a P515/535 emitter-detector module (Walz). Chlamydomonas cells were dark-adapted for 1 h under continuous shaking (120 rpm), and an ECS signal was recorded to study the induction and relaxation of thylakoid membrane energization. And then same cultures were preilluminated (50 μmol photons m^-2^ s^-1^) for 5 min and followed by 4 min dark to study light-induced relaxation kinetics of the ECS signal at 515 nm.

## Results

### Osmotic stress affects the growth and morphology of *C. reinhardtii* under the higher concentration of PEG

In photosynthetic organisms, environmental stress mostly leads to generating reactive oxygen species (ROS) (Laloi et al., 2004; Nagy et al., 2018). Under mild stress conditions, photosynthetic organisms regulate ROS production by increasing ROS scavenging activity (Laloi et al., 2004), induction of NPQ (Erickson et al., 2015), and upregulation of alternative electron transport pathways (Erickson et al., 2015). This first line of defence can be termed acclimation, where little or no changes are observed in growth and cell morphology. If the stress is severe and the acclimation process cannot provide protection, the *C. reinhardtii* cells induce a second line of defence mechanisms ranging from forming multicellular structures to programmed cell death (de Carpentier et al., 2019).

Under mild osmotic stress (2% PEG w/v), the growth, structure and size of *C. reinhardtii* cells were similar to that of control cells (0% PEG w/v). Whereas increased PEG concentrations (5% and 7.5% PEG w/v) reduced growth, cell size, and palmelloid structures (Figure1A). The cell motility is not affected up to 2% PEG concentrations, whereas motility is completely inhibited at higher concentrations of PEG (5% and 7.5%). It is evident from this observation that osmotic stress affects the growth and morphology of *C. reinhardtii* at higher concentrations of PEG. This is one reason we chose to continue our further investigations with higher PEG concentrations in media (7.5% PEG w/v).

### Photosynthetic performance is negatively affected under osmotic stress

Photosynthetic efficiency (F_v_/F_m_) is negatively affected by PEG-induced osmotic stress (Figure 1C). To study the changes in the physiological status of photosynthetic apparatus under osmotic stress, we measured the basic photosynthetic parameters using rapid light curves (RLC) (Figure 1B). Photosynthetic parameters like electron transport rate of PSII ETR(II), photochemical quenching (qP), constitutive energy loss Y(NO) and quantum yield of regulated thermal energy dissipation Y(NPQ) not significantly affected under low PEG concentrations. A significant effect in these parameters is observed under higher concentrations of PEG (7.5%). Under osmotic stress (PEG 7.5%), down-regulation of the electron transport rate of PSII (ETR (II)), the proportion of open reaction centers (qP).

**Figure 1.**
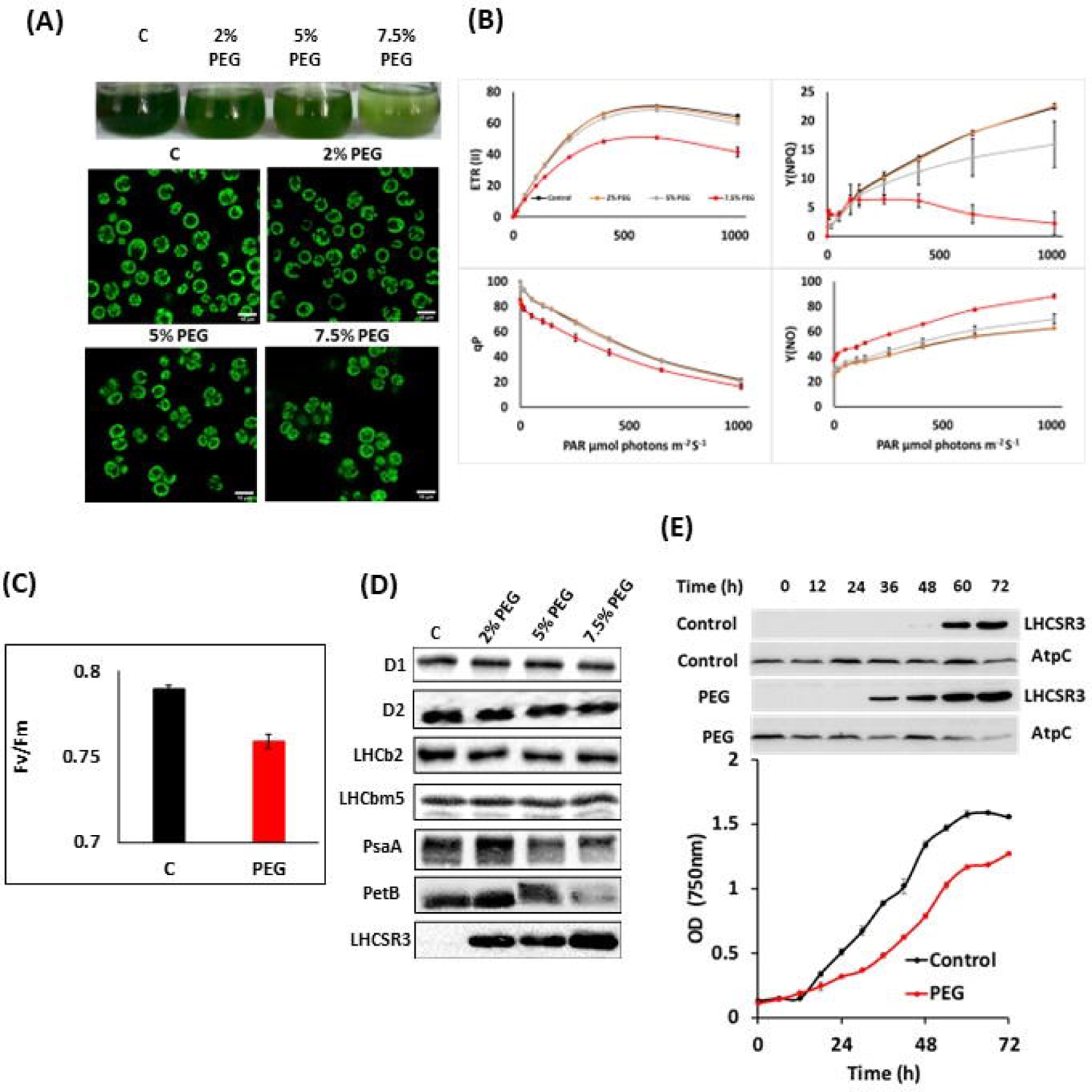
Osmotic stress reduced the growth and photosynthetic efficiency of *C. reinhardtii*. It also leads to the expression of LHCSR3 under all concentrations of PEG. A) Erlenmeyer flask showing growth of *C. reinhardtii* cells grown in under different concentrations of PEG for 72 h (Control-C 0% PEG; 2% PEG, 5% PEG and 7.5% PEG) under continuous illumination of 50 μmol photons m^−2^ s^−1^ (Upper panel) and confocal microscopic image of the same cells (lower panel). B) Light response curves showing electron transport rate of PSII (ETR II), yield of non-photochemical quenching Y(NPQ), photochemical quenching (qP) and non-regulated non-photochemical energy loss Y(NO). Values represent means ± SD (n = 3). C) Photochemical efficiency Fv/Fm of the *C. reinhardtii* cells grown under control (C) and osmotically stressed conditions (7.5% w/v PEG). Values represent means ± SD (n = 3). D) Immunoblot analysis of total cellular proteins from cells grown in the medium supplemented with different concentrations of PEG. The protein levels of D1, D2, LHCb2, LHCBM5 PsaA, PetB, and LHCSR3 can be seen. E) *C. reinhardtii* growth phases with LHCSR3 expression from control (C) and osmotically stressed cells (7.5% w/v PEG), AtpC is used as loading control.

Whereas Y (NO), constitutive energy loss due to non-regulated energy dissipation remained higher than control cells throughout different light intensities. These high values of Y(NO) with the low value of Y(NPQ) indicate an altered physiological state of photosynthetic apparatus inefficient photoprotection by NPQ processes. Therefore, compromised photosynthetic efficiency (Figure 1C) under osmotic stress could be due to the alteration in the physiological state of the photosynthetic apparatus. As changes in photosynthetic parameters and growth alterations were observed under the higher concentration of PEG, we chose to continue our further investigations with higher PEG concentrations in media (7.5% PEG w/v) which will be denoted as “PEG” in rest of the manuscript.

### Analysis of protein content involved in photosynthesis reveals that osmotic stress can induce LHCSR3 expression

To determine the changes in various proteins associated with photosynthetic membranes, the cells collected from different concentrations of PEG and performed immunoblot analysis of specific proteins for PSII reaction center (D1 and D2), major LHCII (LHCb2), minor LHCII (LHCBM5), PSI reaction center PsaA and stress related LHCSR3 (Figure 1D). The immunoblot analysis of PSII reaction center proteins, D1 and D2 has not shown any signs of damage along with their associated LHC’s at any given concentrations of PEG. Whereas, significant decline in the PSI reaction center protein PsaA and Cyt b_6_f complex protein PetB can be clearly observed under higher concentrations of PEG (7.5% w/v).

Under abiotic stress, photosynthetic organisms recruit several unique proteins to alleviate damage to the photosynthetic apparatus. LHCSR3 is one of the proteins expressed under various photooxidizing conditions. It is a well-established phenomenon that LHCSR3 is expressed under high light (Peers et al., 2009) and specific nutrient starvations like iron, copper, sulfur and phosphorous (Im et al., 2003; Zhang et al., 2004; Moseley et al., 2006; Naumann et al., 2007; Strenkert et al., 2016). Here we checked whether LHCSR3 could be expressed under osmotic stress. The immunoblot analysis of proteins isolated from *C. reinhardtii* cells grown under various concentrations of PEG indicated that LHCSR3 is expressed, and its expression levels are high at higher concentrations of PEG in the medium (Figure 1D). Therefore, LHCSR3 expression is possible under osmotic stress and does not require HL.

Nutrient limitation can stunt the growth of organisms, and it can be studied by growth curve analysis (Figure 1E). In the batch culture of *C. reinhardtii* grown under different media compositions (with PEG (PEG) and without PEG (Control)), the stationary phase reached after 60h of growth. In the cells grown under control conditions, LHCSR3 expression is observed at the stationary phase (60 h), whereas, under osmotic stress, the growth rate is low, and LHCSR3 expression is observed at the mid logarithmic phase of growth (36 h) (Figure 1E). Therefore, either diminished growth rate under osmotic stress or stationary phase in control cells can express LHCSR3 protein.

### Photooxidizing conditions are necessary for LHCSR3 expression under osmotic stress

In *C. reinhardtii,* LHCSR3 expression is observed when the cells are either grown under high light (Peers et al., 2009) or under nutrient-deficient conditions (Zhang et al., 2004; Moseley et al., 2006; Naumann et al., 2007; Strenkert et al., 2016). Moreover, in several studies with nutrient starvation, LHCSR3 expression is observed when grown under moderate light intensity (Zhang et al., 2004; Moseley et al., 2006; Naumann et al., 2007; Strenkert et al., 2016). As mentioned above, LHCSR3 expression occurs in various light intensities; we are interested to know the role of light in its expression under osmotic stress. To answer this question, *C. reinhardtii* cells were grown in a TAP medium under four different conditions for 72h. The first set of cells are control cells grown under growth light (no stress), the second set of cells in PEG-supplemented TAP medium under growth light (osmotic stress and optimum light intensity), and the third set in TAP medium under high light, HL (optimum medium composition and high light stress) and fourth set of PEG supplemented TAP medium under dark (osmotic stress with heterotrophic growth condition). The immunoblot analysis from the above samples indicates that LHCSR3 expression is not observed in the control cells, dark-grown cells, and cells grown in PEG supplemented medium under dark (Figure 2A). Whereas its expression is clearly seen in cells grown in PEG supplemented medium under growth light conditions and in HL. Therefore, osmotic stress alone cannot induce LHCSR3 expression; thus, LHCSR3 expression requires photo-oxidative stress where photosynthetic electron transport and other light induced signaling pathways are necessary for LHCSR3 expression. Hence, we conclude that light is essential for expressing LHCSR3 under osmotic stress.

**Figure 2.**
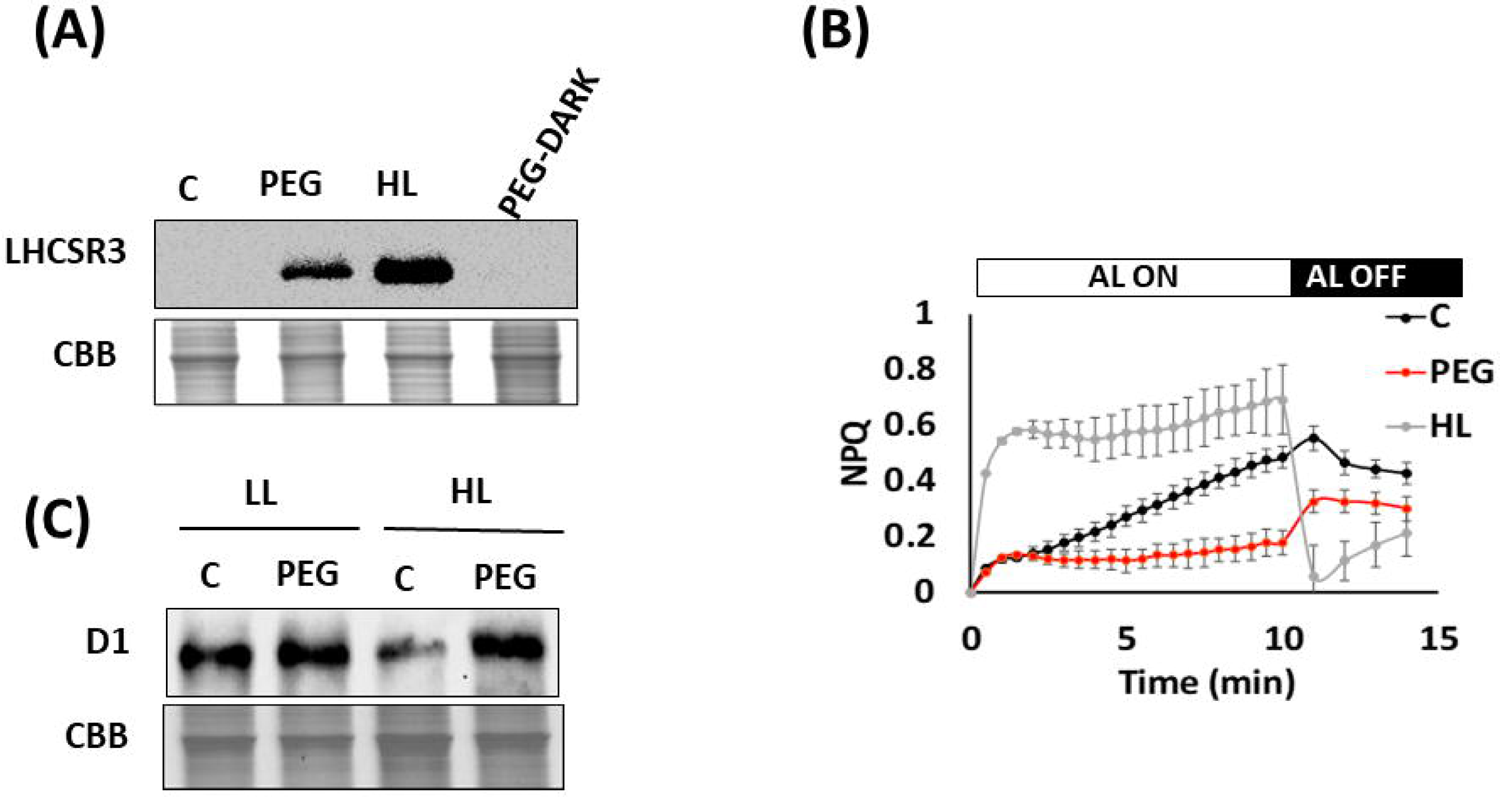
Photooxidizing conditions are necessary for LHCSR3 expression under osmotic stress, and its photoprotective function is not dependent on qE. A) LHCSR3 expression in the cells grown under different conditions (control) C (TAP medium at 50 μmol photons m^−2^ s^−1^ of illumination), PEG (TAP medium supplemented with 7.5% w/v of PEG at 50 μmol photons m^−2^ s^−1^ of illumination), HL (in TAP medium at 250 μmol photons m^−2^ s^−1^ of illumination) and PEG-dark (in TAP medium at 0 μmol photons m^−2^ s^−1^ of illumination). B) NPQ of cells grown in different conditions. Values represent means ± SD (n = 3). C) Immunoblot analysis for quantifying D1 protein content after incubation in chloramphenicol (CAM) 100 µg/ml.

### NPQ component qE is not active despite LHCSR3 expression under osmotic stress

Figure 2B shows NPQ induction kinetics in the *C. reinhardtii* cells grown photoheterotrophically (TAP medium) under growth light (∼50 μmol photons m^-2^ s^-1^) (control cells), HL (∼500 μmol photons m^-2^ s^-1^) and PEG supplemented TAP medium (osmotic stress). As expected, NPQ induction and relaxation are fast in HL-grown cells due to the expression of LHCSR3 protein (Figure 2B). Here, most NPQ is dominated by the qE component. Whereas, in control cells and osmotically stressed cells (PEG), NPQ induction and relaxation are slow, indicating the involvement of other components of NPQ except for qE. In control cells, a slow rise in NPQ after some minutes suggests the participation of qZ and/or qT upon dark to light transition. The absence of a slow increase in NPQ of PEG-grown cells could be attributed to hampered NPQ mechanisms like state transitions under osmotic stress (Figure S1). These results suggest that the impairment of known NPQ mechanisms under osmotic stress and LHCSR3 expression does not result in qE upon exposure to high light.

### LHCSR3 alleviate D1 damage under osmotic stress

The above data shows that LHCSR3 protein is expressed in the cells grown under osmotic stress. Of all the proteins involved in photosynthetic electron flow, PSII reaction center protein D1 is more susceptible to damage under photooxidizing conditions. LHCSR3 being a strong quencher of excitation energy, is expected to protect PSII reaction center protein D1 under photo-oxidative stress. Visualization of D1 damage is tricky due to the high turnover rate of D1 protein. To observe D1 damage its de novo synthesis must be inhibited by chloroplast protein synthesis inhibitor like chloramphenicol. In order to establish the photoprotective role of LHCSR3 expression under osmotic stress, control cells grown under moderate light intensity (50 µmol photons m^-2^ s^-1^) which do not have LHCSR3 expressed and PEG-grown cells same light conditions with LHCSR3 expressed are treated with chloramphenicol (Kuroda et al., 2014) in the dark for 30 min (chloroplast protein synthesis inhibitor), and these cells are exposed to HL (500 μmol photons m^-2^ s^-1^) for 2 h. The immunoblot analysis of total cellular proteins from these cells shows that under HL, D1 protein is not damaged in PEG-grown cells. In contrast, control cells exposed to HL suffered extensive D1 damage (Figure 2C). This experiment clearly shows that the expression of LHCSR3 under osmotic stress alleviates D1 damage and plays an essential role in the photoprotection of PSII.

### LHCSR3 expression prevents ROS formation under osmotic stress

Over-excitation of chlorophylls or imbalance between photoreception and utilization can result in ROS. Comparing ROS levels in WT and *npq4* strains grown under osmotic stress clearly shows the higher ROS levels in *npq4* mutant under osmotic stress (Figures 3A, B). This indicates the role of LHCSR3 in preventing ROS formation.

**Figure 3.**
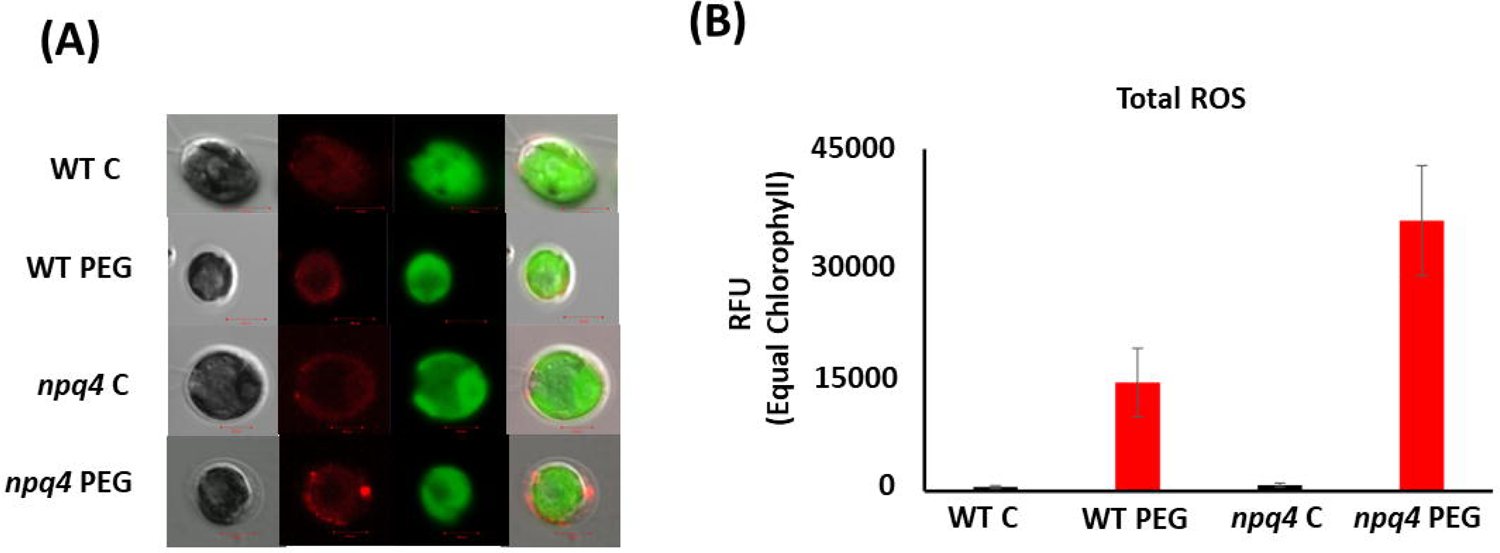
LHCSR3 expression reduces ROS production under osmotic stress. A) Confocal microscopy images for the ROS content in the cells grown under osmotic stress. WT and *npq4* strains are grown under control (C) and osmotic stress (PEG) conditions, and images were captured using 2,7dichlorodihydrofluorescein diacetate (H_2_DCFDA) (10 μM) staining. B) Total ROS in the cells was quantified using spectroscopy at equal chlorophyll concentration. Values represent means ± SD (n = 3).

### Significant structural changes in the thylakoid membrane are observed under osmotic stress

The thylakoid membrane is dynamic. It can undergo several structural changes to acclimate to environmental stress (Nagy et al., 2014). Here, we want to know the impact of LHCSR3 expression on thylakoid membrane architecture and the supramolecular organisation of protein complexes. We grew *C. reinhardtii* cells wild type and *npq4* mutant (that lacks LHCSR3 gene expression) in the presence and absence of osmotic stress. Changes in thylakoid membrane structure and composition were analysed using CD spectroscopy and lpBN-PAGE.

The visible red region of the CD spectrum of *C. reinhardtii* cells consists of two intense *psi*-type bands at (-) 675 and (+) 690 nm (Nagy et al., 2014). These *psi*-type bands are given rise by the presence of large ordered arrays (size ranging from 200 to 400 nm) of PSII-LHCII complexes in the appressed region of the thylakoid membrane. The magnitude of the *psi*-type band is proportional to the size of the ordered array. In WT *C. reinhardtii* cells grown under osmotic stress, there is a significant decrease in the amplitude of the red *psi*-type signal, whereas we observed marginal changes in the *psi*-type signal from *npq4* mutant grown under osmotic stress (Figure 4A). This indicates that LHCSR3 expression affects thylakoid structure under osmotic stress. Moreover, the stronger *psi*-type signal from WT cells grown under optimum conditions (control) indicates that the macroarrays composed of PSII-LHCII supercomplexes are larger in control cells than in PEG-grown cells (Garab and van Amerongen, 2009; Nagy et al., 2014). The excitonic signal (-) 650 nm rises from the trimeric array of chlorophyll *b* complexed with LHCII. In WT cells grown under osmotic stress, the decreased amplitude of excitonic CD signal (-) 650 nm compared with *npq4* mutant indicates an alteration in LHCII trimeric structure. Therefore, osmotic stress effects the PSII-LHCII super-complex structure and could be facilitated by LHCSR3 expression.

**Figure 4.**
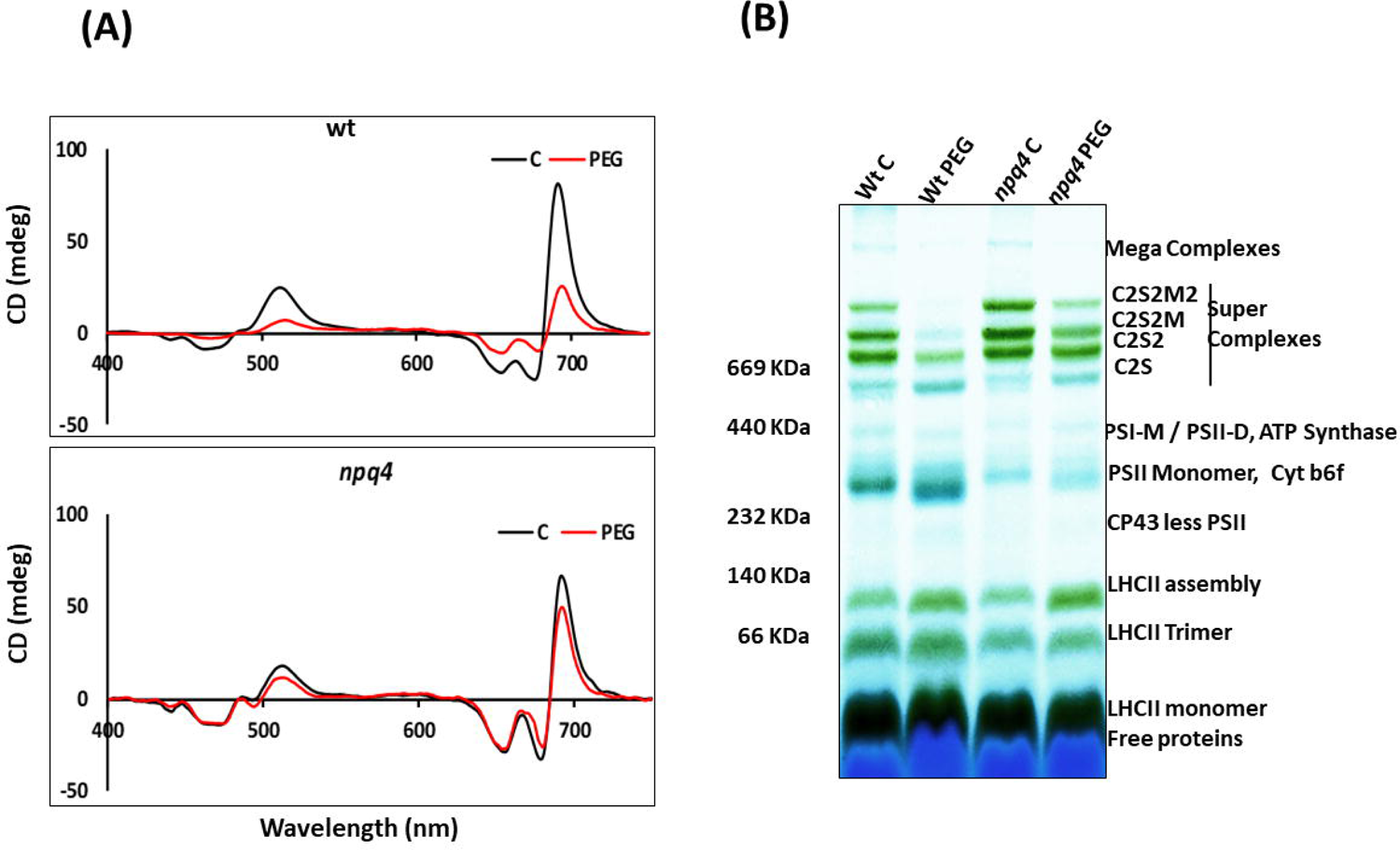
LHCSR3 expression is associated with changes in thylakoid membrane architecture under osmotic stress. A) CD analysis of the cells grown under osmotic stress. The upper panel shows the changes in thylakoid macrodomain assembly in wild type cells and the lower panel show changes in *npq4* mutant. B) lpBN-PAGE of the thylakoid membranes from WT and *npq4* mutant grown in (control) C and (osmotic stress) PEG. Thylakoid membranes were solubilized in 1% βDM.

Thylakoid membranes comprise several multiprotein membrane complexes that run photosynthetic light reactions. Their composition, distribution, and stoichiometry may vary according to the changes in the environmental conditions. Thylakoids are composed of three structures: grana, stroma and grana margins. The appressed regions of thylakoids, grana, are enriched with PSII-LHCII supercomplexes, and non-appressed regions, stroma, with PSI-LHCI and ATPase complexes (Nevo et al., 2012). Grana margins have been suggested to have both photosystems PS (I and II) that can form mega complexes (Järvi et al., 2011). The localization of the Cyt b_6_f complex is under debate, but many studies reported its even distribution in the thylakoid membrane (Anderson, 1982; Albertsson et al., 1991). To study the composition and dynamics of thylakoid membrane pigment-protein complexes under osmotic stress, *C. reinhardtii* WT and *npq4* mutant were grown in TAP medium supplemented with PEG (Osmotic stress)/without PEG (control) and thylakoids were isolated. Prior to native-PAGE, proteins are isolated from the thylakoid membrane using mild non-ionic detergent, β-dodecyl maltoside (β-DM) (1% final concentration). We opted to use β-DM for its ability to solubilize most of the thylakoid membrane, but it comes at the expense of losing weak hydrophobic interactions in protein complexes (Wittig et al., 2006). Preliminary identification of these protein complexes was performed by immunoblotting 1D of lpBN-PAGE with antibodies specific for PSII RC protein D1, PSI RC protein PsaA, and Cyt b_6_f complex protein PetA (Figure S2 *A*) and bands were named according to previous studies (Rantala et al., 2017; Rantala et al., 2018).

Figure 4B show typical patterns of thylakoid membrane protein complexes separated in lpBN-PAGE. Four bands of PSII-LHCII supercomplexes near and above 669 kDa, PSII and PSI monomers devoid of LHCs near 440 kDa and lower three bands consisting of different hierarchal combinations of LHC have been resolved from the thylakoids of *C. reinhardtii* (WT control and *npq4* control) cells grown under optimum conditions. The largest PSII supercomplexes isolated from *C. reinhardtii* have two reaction centres attached with six LHC trimers (C_2_S_2_M_2_L_2_) (Burton-Smith et al., 2019). LHCII trimers are named as strongly bound trimers (S-trimer), moderately bound trimers (M-trimer), and loosely bound trimers (L-trimer), depending on their binding strength with PSII core complex (C). Here, in lpBN-PAGE, the four high molecular weight bands (around (669 kDa) consist of different combinations of PSII core with S and M trimers, and the largest supercomplex observed is C_2_S_2_M_2_, as interactions with L-trimers are lost during solubilisation with (β-DM) (Wittig et al., 2006).

Analysis of lpBN-PAGE of thylakoids isolated from the cells grown under osmotic stress shows the decreased intensity of the PSII-LHCII supercomplex in WT cells. In contrast, the PSII-LHCII supercomplexes’ intensity is less affected in the *npq4* mutant (Figure 4B). The dissociation of LHCII trimers from the PSII core resulted in increased intensity of the C_2_S, PSII Monomer and LHCII populations that migrated to lower molecular weight regions. In *npq4* mutant, these changes are less pronounced.

Immunoblot analysis from the second dimension of lpBN-PAGE of thylakoids from cells grown under osmotic stress (Figure S2*B*) shows that LHCSR3 migration is associated with either LHCII assembly or monomeric LHC. Therefore, destabilization of the PSII-LHCII complex could be related to LHCSR3 expression under osmotic stress, and the observation confirms that this change is insignificant in the *npq4* mutant.

### Downregulation of chloroplast ATP synthase content under osmotic stress is responsible for the expression of LHCSR3

Thylakoid lumen acidification is significant for the function and expression of LHCSR3. The accumulation of protons in the thylakoid lumen generates membrane potential (Δψ) and proton gradient (ΔpH) across the thylakoid membrane. This Δψ and ΔpH contribute to the proton motive force (*pmf*). This *pmf* drives protons through (cp) ATP synthase. Thylakoid lumen acidification is achieved when the rate of proton translocation into the thylakoid lumen is more than the rate of H^+^ conductance by (cp) ATP synthase. This generally occurs under high light due to the increased rate of photosynthetic electron flow.

Changes in membrane potential of thylakoid membrane can be observed by the electrochromic shift (ECS P515) signal and this could be the poxy measurement for thylakoid lumen acidification. Figure.5A shows single turnover flash-induced changes in ECS signal from *C. reinhardtii* cells. P515 signal measures electric potential across thylakoid membrane due to electrochromic pigment absorption of carotenoids and Chl *b* (Duysens, 1954, Junge and Witt, 1968). The study of normalized ECS signal recorded for both control and PEG-grown cells shows the rapid increase in signal due to primary charge separation in PSII and PSI reaction centres (Figure 5A) (upper panel). This rapid rise in signal is less in PEG-grown cells than in the control. Moreover, in control cells, this rapid increase in signal is followed by a slower decay (t_1/2_ approx. 20 ms) due to the electrogenic Q-cycle at Cyt b_6_f complex (Velthuys, 1978). This slow decay in signal is less pronounced in PEG-grown cells, which could be possibly attributed to the low content of functional Cyt b_6_f complex (Figure 1D) and the low electron transport rate of PSII (Figure 1B).

**Figure 5.**
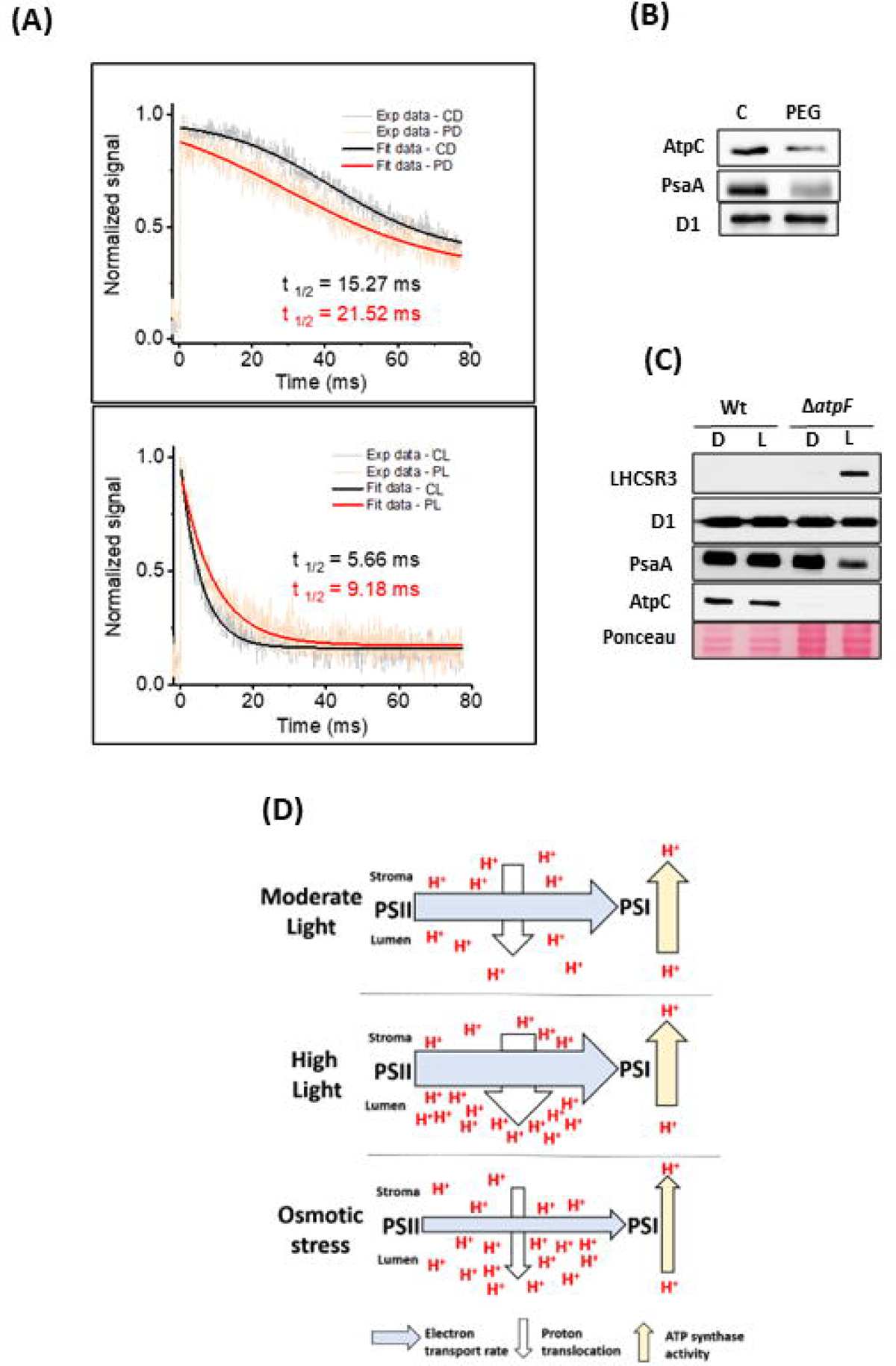
Under osmotic stress, reduced ETRII is compensated by downregulation of (cp) ATP synthase complex to generate enough. Δ**pH required for LHCSR3 expression.** A) Normalized P515 absorption kinetics of the *C. reinhardtii* cells grown under PEG induced osmotic stress. CD - control cells grown without PEG and dark adapted for 1 h; PD-osmotically stressed cells grown with 7.5% PEG dark adapted for 1 h; CL control cells exposed to light for 5 min; PL osmotically stressed cells exposed to light for 5 min B) Immunoblot analysis of changes involved in ATP synthase content under PEG induced osmotic stress. C) Expression of LHCSR3 in (cp)ATP synthase mutant (*atpF)* when exposed to light. D) Model depicting the possible mechanism for the development of ΔpH under osmotic stress. The electron transport rate is optimum under moderate light intensity; here, the proton translocation from stroma to thylakoid lumen is balanced by (cp)ATP synthase activity that utilizes ΔpH that can prevent lumen acidification (top panel). Under highlight, the increased electron transport rate thus increased proton translocation into thylakoid lumen, is not balanced by (cp)ATP synthase activity resulting in thylakoid lumen acidification (middle panel). Under osmotic stress with moderate light intensity, the decreased rate of electron transport and compromised (cp)ATP synthase activity results in thylakoid lumen acidification as the latter mechanism is more affected (bottom panel). The Width of the arrows represents the comparative magnitude of that specific activity in each growth condition.

The decay of the P515 signal is highly dependent on the physiological status of the sample. This decay occurs specifically due to H^+^-ion conductance by (cp) ATP synthase that requires illumination. To study the ATP synthase activity, *C. reinhardtii* cells were pre-illuminated with actinic light (50 μmol photons m^-2^ s^-1^) (to activate ATP synthase) for 5 min and dark-adapted for 4 min before recording rapid P515 induction/ relaxation kinetics by single turnover flash. Figure. 5A (lower panel) shows the rapid P515 relaxation kinetics of the *C. reinhardtii* cells grown in the presence or absence of PEG. Fast relaxation of P515 signal from control cells than PEG grown cells indicates the low turnover rate of ATP synthase complex under osmotic stress. The immunoblot analysis of thylakoids isolated from control and PEG grown cells also show the downregulation of ATP synthase content in PEG grown cells (Figure 5B). This effect could be observed in slow relaxation of the proton gradient under osmotic stress (Figure 5A). Moreover, there is low abundance of PsaA protein in osmotically stressed cells (Figure 5B) and atpF mutant when exposed to light (Figure 5C), indicating that PSI is sensitive to light when ATP synthase activity is affected.

Previous studies on higher plants indicate the relationship between the downregulation of ATP synthase activity and high *pmf* (Kohzuma et al., 2009; Davis et al., 2016). *pmf* is stored in the form of two components Δψ and ΔpH. Increased ΔpH a component of *pmf* can induce energy-dependent quenching qE by protonating PSII subunit PsbS (Takizawa et al., 2007) and LHCSR3. Therefore, the downregulation of ATP synthase activity can induce NPQ through ΔpH. In *C. reinhardtii,* qE depends on the expression of LHCSR3, whereas LHCSR3 expression and function depend on thylakoid lumen acidification. Further, thylakoid lumen acidification is regulated by ATP synthase activity. To establish the relationship between ATP synthase down-regulation and LHCSR3 expression, *C. reinhardtii atpF* mutant (CC-4150) deficient in CF0 of ATP synthase complex (Bennoun et al., 1980) is grown in a TAP medium under low light (∼5 μmol photons m^-2^ s^-1^) and then exposed to moderate growth light (30 μmol photons m^-2^ s^-1^) for 2 h. The immunoblot analysis of total cellular proteins from *atpF* mutant exposed to light showed the expression of LHCSR3 protein (Figure 5C). Therefore, these results indicate that in *C. reinhardtii*, there is a significant relationship between LHCSR3 expression and the downregulation of ATP synthase content under osmotic stress.

## Discussion

Under saturating light conditions, the increased photosynthetic electron flow can lead to an imbalance between light energy absorption and its utilization. This imbalance can result in feedback regulation of photosynthesis through NPQ (Müller et al., 2001). In *C. reinhardtii,* LHCSR3 expression is required for induction of qE (Peers et al., 2009), an essential component of NPQ and its expression is observed in various photooxidizing conditions like high light (Peers et al., 2009), blue light (Petroutsos et al., 2016), UV light (Allorent et al., 2016), and nutrient starvation (Im et al., 2003; Zhang et al., 2004; Moseley et al., 2006; Naumann et al., 2007; Strenkert et al., 2016). The mechanism of LHCSR3 expression is well studied under HL, and these studies emphasized the importance of thylakoid lumen acidification due to increased photosynthetic electron transport rate. The direct link between thylakoid lumen acidification and induction of LHCSR3 expression is not clearly established. However, the presence of Ca^2+^/ H^+^ antiporter (Ettinger et al., 1999) for light-driven uptake of calcium by thylakoid lumen and dependence of LHCSR3 expression on the CAS (Petroutsos et al., 2011) signalling system indicate the possible role of ΔpH in signalling. Whereas, under specific nutrient starvations, where LHCSR3 expression is reported, downregulation of the photosynthetic electron transport rate is also observed (Zhang et al., 2004; Moseley et al., 2006; Terauchi et al., 2010). This leads to some fundamental questions that how ΔpH is achieved under reduced electron transport rate and the function of LHCSR3 expressed under nutrient limitation.

Osmotic stress limits water uptake by plants and cell cultures. Most of the nutrients absorbed by plants are water-soluble; therefore, osmotic stress could be called nutrient starvation. We choose PEG-8000 as an osmoticum for our experiments due to specific properties like its large size, osmotic potential, inert nature and negligible penetrability into the cells. Here, we compared the growth characteristics, morphological changes and photosynthetic performance of the *C. reinhardtii* cells grown in a PEG supplemented medium with previous reports of NaCl-induced osmotic stress (Khona et al., 2016). Reduction of growth rate (Figure 1E), cell size, palmelloid formation (Figure 1A) and decreased photosynthetic performance (Figure 1B) are clear indications of osmotic stress at higher concentrations of PEG.

### Photoprotective role of increased constitutive energy loss Y(NO) under osmotic stress

Optimum photosynthesis requires delicate coordination between the absorption of light energy, carbon fixation and regulation of photosynthetic activity. This delicate balance is altered by several environmental stress factors that can result in low photosynthetic yield; moreover, under severe stress conditions, it leads to photoinhibition. The down-regulation of electron transport rate ETR (II), photochemical energy utilization qP and regulated dissipation of excess energy Y(NPQ) in PEG-grown cells (Osmotic stress) is the indication of a negative impact on photosynthetic performance (Figure 1B). Whereas, under osmotic stress, the increase in constitutive energy loss, also called non-regulated energy quenching Y (NO) (Hendrickson et al., 2004; Kramer et al., 2004), indicates the inability of PSII to utilize the incident radiation for photosynthesis. Here, in this case, Y(NO) remained high under increasing light intensity (Figure 1B) and with constant saturating light intensity (Figure S3A). This, constitutive thermal dissipation (Y (NO)) can happen due to damage to the PSII and/or the inability of the other photosynthetic complexes like Cyt b_6_f complex and PSI to cope up with the rate of electron flow from PSII. To understand this phenomenon under osmotic stress, immunoblot analysis is performed for the proteins involved in photoreception and electron transport chain (Figure 1D). It is interesting to note that D1 protein, known to be sensitive to photo-oxidative stress (Nishiyama et al., 2001), is stable under different concentrations of PEG. The same is true for some of its LHCII antenna proteins.

The protein contents of photosynthetic membrane complexes downstream of ETC of PSII like PetB and PsaA are down-regulated under higher concentrations of PEG. As discussed earlier, osmotic stress limits water uptake by cells, thus, less nutrient permeability into the cells. The loss of PSI and Cyt b_6_f components under osmotic stress could be attributed to the unavailability of certain metal ions required for their synthesis (Terauchi et al., 2010). Moreover, the movement of CO_2_ into Chlamydomonas cells requires water. CO_2_ in its soluble form enters the Chlamydomonas cells with the aid of HCO^3−^ transporters which can be converted back to CO_2_ by carbonic anhydrase (Moroney et al., 2011). Less water uptake under osmotic stress also affects inorganic carbon availability. This can cause an imbalance between the photoreception of PSI and carbon fixation reactions that could result in PSI photoinhibition (Nishiyama et al., 2001). Therefore, increased constitutive energy loss from PSII could be the mechanism to balance the loss of the Cyt b_6_f complex and PSI under extreme photo oxidizing conditions. This strategy could prevent the over-reduction of inter-system electron carriers under severe environmental stress (Bag et al., 2020). This increased Y(NO) phenomenon without change in D1 protein content indicates its photoprotective role when NPQ systems are compromised under extreme environmental stress (Barbato et al., 2020).

### Alternative role of LHCSR3 under osmotic stress

LHCSR3 is a strong quencher of excitation energy under saturating light intensities. In *C. reinhardtii* cells grown under HL, LHCSR3 expression is responsible for the induction of fast and reversible components of NPQ (Peers et al., 2009). In contrast, its function under other abiotic stress conditions is not clear. Interestingly, under osmotic stress, LHCSR3 expression can occur at moderate light intensity (Figures 1D, 2A), with no observable fast and reversible component of NPQ (Figure 2B). To understand if this LHCSR3 can protect PSII under HL, immunoblot analysis for quantification of D1 protein was performed by blocking *de-novo* proteins synthesis with chloramphenicol (D1 protein has high turnover) and exposing the cells to HL for 2h. Under HL, D1 damage is observed in control cells, and there is no apparent reduction of D1 content in PEG-grown cells (Figure 2C). In the above experimental conditions, LHCSR3 expression is only observed under PEG-induced osmotic stress (Figure 2A), suggesting LHCSR3-mediated photoprotection of PSII. Moreover, the study of ROS generation with WT and *npq4* strains grown under osmotic stress indicates the role of LHCSR3 in preventing ROS under osmotic stress (Figures 3A,B).

These results raise the question that how LHCSR3 achieves photoprotection of PSII without induction of qE? To answer this question, we studied the changes in thylakoid membrane architecture using non-invasive CD spectra of intact cells and changes in the supramolecular organisation of thylakoid protein complexes using lpBN-PAGE. In WT cells where LHCSR3 is expressed under osmotic stress, the thylakoid membrane architecture and composition of PSII-LHCII super complex are affected. The weaker *psi*-type signal in WT cells grown under osmotic stress indicates major rearrangement in the macromolecular arrays of PSII-LHCII complex enriched grana. Whereas in *npq4* mutant that lacks LHCSR3 gene expression, show little change in thylakoid remodelling under osmotic stress (Figure 4A). The same can be observed from the composition of thylakoid membrane protein complexes in lpBN-PAGE analysis (Figure 4B). In *C. reinhardtii,* PSII-LHCII comprises two reaction center cores and six LHCII trimers C_2_S_2_M_2_L_2_ (Burton-Smith et al., 2019). Here, in this lpBN-PAGE, due to solubilisation with β-DM, L-trimers dissociate from PSII-LHCII supercomplexes leaving different combinations of the core with S and M trimers (Wittig et al., 2006). In WT cells grown under osmotic stress, less abundance of PSII-LHCII complexes is observed, whereas, in *npq4* mutant, this change is less pronounced. The role of other molecular players that partly compensate for the LHCSR3 function in thylakoid remodelling under osmotic stress needs to be investigated. This result from lpBN-PAGE complements with the results from CD spectra, where higher-order structures in the thylakoid membrane are affected under osmotic stress. In support of the above discussion, previous studies using single particle analysis of PSII-LHCII-LHCSR3 (C_2_S_2_-LHCSR3) complex isolated (using α-DM) from wild-type *C. reinhardtii* (137c) cells grown under HL did not show any bound M and L trimers. In the npq4 mutant, the largest particles isolated consist of C_2_S_2_M and C_2_SML (Semchonok et al., 2017). The same phenomenon could be a reason for the stronger intensity of higher order PSII-LHCII complexes in *npq4* mutant (Figure 4B). In another study using the same detergent showed the association of few LHCSR3 protein with PSII-LHCII complex (Tokutsu and Minagawa, 2013); however, most of the LHCSR3 proteins are observed either with higher order LHCs (LHC II trimers) or with LHCII monomers. A similar result has been observed in immunoblot analysis of lpBN-2D PAGE of thylakoids isolated from WT cells grown under osmotic stress, where LHCSR3 is association/ migration with higher order LHCII and LHCII monomers (Figure S2B). These observations suggest that the expression of LHCSR3 under osmotic stress could weaken the interactions between the PSII core and the LHC II complex. Thus, preventing the transfer of excitation energy to the most vulnerable region of the PSII core. Here we propose a possible mechanism of LHCSR3 mediated thylakoid membrane remodelling that involves dissociating LHCII from PSII to confer photoprotection under osmotic stress (Figure 6).

**Figure 6.**
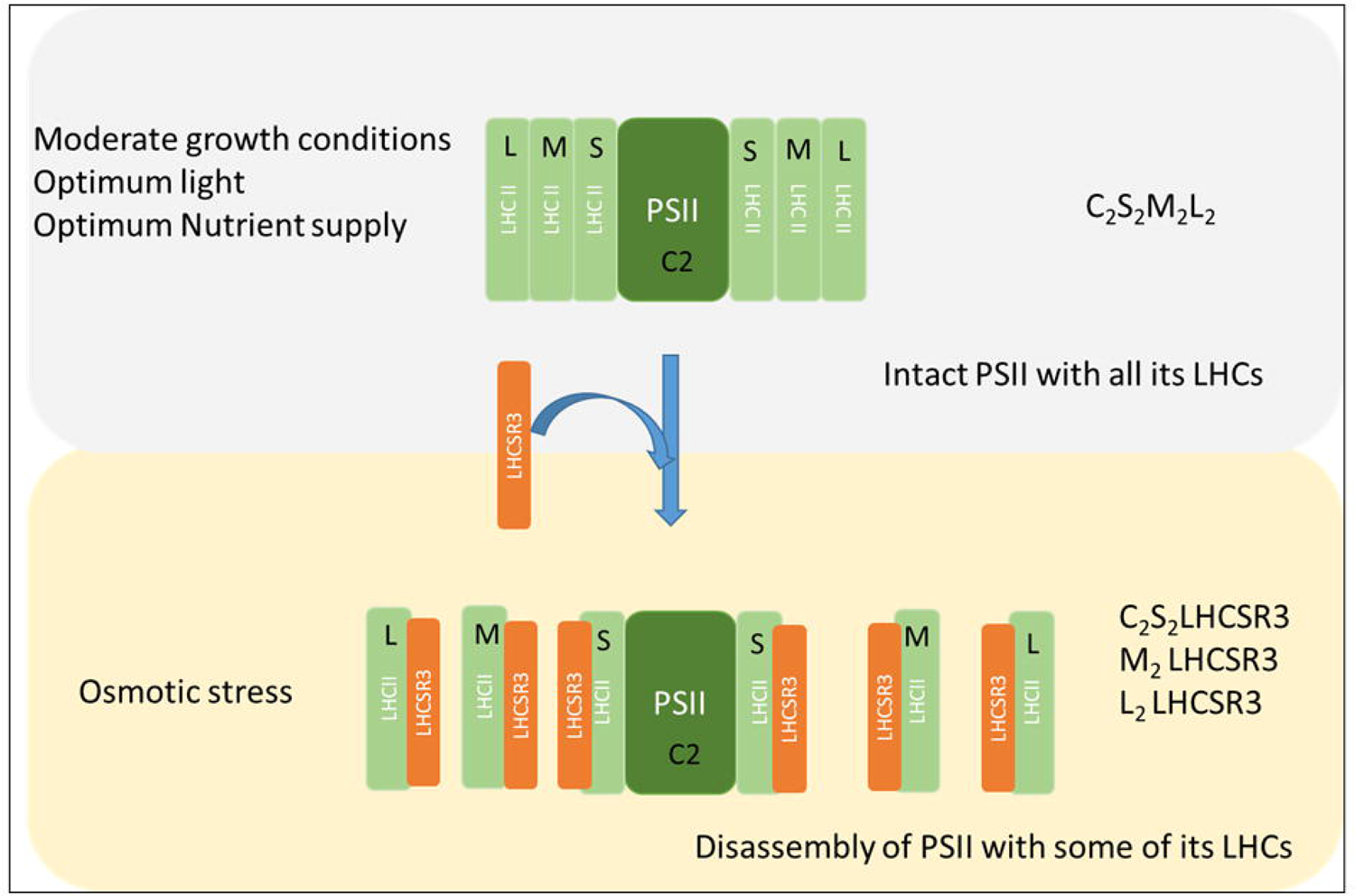
Proposed model to indicate that LHCSR3 confers photoprotection under osmotic stress. In *C. reinhardtii,* under moderate light conditions and nutrient supply, the PSII supercomplex exists as the C_2_S_2_M_2_L_2_ complex. Under photooxidative stress like osmotic stress, the recruitment of LHCSR3 may facilitate the dissociation of most of the LHC II molecules attached to the PSII core and excitation energy transfer from LHCII to PSII core could be inhibited, preventing photooxidative damage to the PSII core.

This antenna detachment model for photoprotection requires a detached LHCII-LHCSR3 complex to function independently as a quenching complex. Recent evidence shows that LHCSR3 is a sensitive pH sensor, and quenching sites are located within LHCs (Tian et al., 2019). Therefore, LHCSR3 and LHCII complexes could function together to quench excitation energy within the complex. Proven sensitivity of LHCSR3 towards the changes in luminal pH (Tian et al., 2019) with slow relaxation of ΔpH due to low ATP synthase activity (Figure 5A, B) and observed sustenance of Chl *a* fluorescence quenching (FLJ and F_m_) (Figure S3B) under osmotic stress in WT and absence of sustanence of quenching in *npq4* mutant suggests the role of LHCSR3 in some form of quenching that can sustain for longer duration. This thylakoid membrane remodelling with LHCSR3 could protect the photosynthetic apparatus (PSII) of *C. reinhardtii* from prolonged unfavourable conditions like nutrient starvation.

### Expression of LHCSR3 under osmotic stress

ΔpH is determined by the balance between photosynthetic electron flow and dissipation of pH gradient by ATP synthase activity (Pinnola and Bassi, 2018). Under HL, increased electron transport results in high proton (H^+^) flux into the thylakoid lumen that can shift luminal pH to acidic. This drop in pH of the thylakoid lumen is sensed by the proton binding residues of LHCSR3 to initiate NPQ within time scale of milliseconds. Another role of the thylakoid lumen might be associated with signal transduction for LHCSR3 expression. LHCSR3 expression requires thylakoid located plant-specific calcium-binding protein (CAS) activity (Petroutsos et al., 2011). Light-dependent Ca^2+^/ H^+^ antiporter in the thylakoid membrane (Ettinger et al., 1999) is the clear indication for the LHCSR3 expression dependent on thylakoid lumen acidification provided by increased ETR under HL. Therefore, decreased electron transport rate (Figure 1B) under osmotic stress raises a fundamental question: how is ΔpH achieved for LHCSR3 expression and function? To answer this question, we studied (cp) ATP synthase content and activity. The reduced ATP synthase content (Figure 5B) and activity (Figure 5A) is observed under osmotic stress in *C. reinhardtii*. Therefore, here we propose that under osmotic stress, low (cp) ATP synthase content and activity results in decreased proton efflux from thylakoid lumen that could result in build-up of ΔpH across thylakoid membrane enough for expression and function of LHCSR3 protein. To verify this hypothesis, we checked the capability of *C. reinhardtii atpF* mutant (CC-4150) impaired in ATP synthase complex assembly (Bennoun et al., 1980; Lemaire and Wollman, 1989) to express LHCSR3 protein when exposed to light. In support of our hypothesis, LHCSR3 expression is clearly observed in this mutant upon exposure to light (Figure 5C). The schematic representation of the mechanism to generate ΔpH under osmotic stress is given in Figure 5D.

In conclusion, LHCSR3 expression under osmotic stress is possible by down-regulation of (cp) ATP synthase content and activity. The qE component of NPQ is not functional despite LHCSR3 expression. Here, we propose an alternative function of LHCSR3 under osmotic stress that LHCSR3 expression prevents the excitation energy transfer from LHCII to PSII RC by weakening interactions between the PSII core and its antenna. Thus, conferring photoprotection to PSII RC and downstream components of ETC. With this work, we suggest that LHCSR3 expression can occur in any condition where the absorbed light energy is not utilised by subsequent metabolic reactions irrespective of light intensity.

## Author Contributions

R.S and S.M designed the research; S.M, R.M.Y, P.B and M.Y.Z performed the research; S.M and R.S analyzed the data; R.S and SM wrote the paper.

## Competing Interest Statement

Authors have no conflicts of interest to declare.

## Acknowledgements

R.S was supported by the Science & Engineering Research Board (CRG/2020/000489), Joint UGC- ISF Research Grant - File No. 6-8/2018 (IC) and Council of Scientific and Industrial Research (No.38 (1504)/21/EMR-ll) Govt. of India, for financial support. DST-FIST and UGC-SAP, Govt. of India, for infrastructure development for the Department of Plant Sciences, University of Hyderabad. SM acknowledged UGC for fellowship (JRF/SRF).SM aknolwdged Justin Findinier for confocal imaging.

## Notes

### Competing Interest Statement

The authors have declared no competing interest.

## References

Albertsson, P.-Å., Andreasson, E., Svensson, P. and Yu, S.-G. (1991) Localization of cytochrome f in the thylakoid membrane: evidence for multiple domains. Biochimica et Biophysica Acta (BBA) - Bioenergetics, 1098, 90–94.

Allorent, G., Lefebvre-Legendre, L., Chappuis, R., Kuntz, M., Truong, T.B., Niyogi, K.K., Ulm, R. and Goldschmidt-Clermont, M. (2016) UV-B photoreceptor-mediated protection of the photosynthetic machinery in Chlamydomonas reinhardtii. Proceedings of the National Academy of Sciences of the United States of America, 113, 14864–14869.

Anderson, J.M. (1982) Distribution of the cytochromes of spinach chloroplasts between the appressed membranes of grana stacks and stroma-exposed thylakoid regions. FEBS letters, 138, 62–66.

Bag, P., Chukhutsina, V., Zhang, Z., Paul, S., Ivanov, A.G., Shutova, T., Croce, R., Holzwarth, A.R. and Jansson, S. (2020) Direct energy transfer from photosystem II to photosystem I confer winter sustainability in Scots Pine. Nat Commun, 11, 6388.

Barbato, R., Tadini, L., Cannata, R., Peracchio, C., Jeran, N., Alboresi, A., Morosinotto, T., Bajwa, A.A., Paakkarinen, V., Suorsa, M., Aro, E.M. and Pesaresi, P. (2020) Higher order photoprotection mutants reveal the importance of ΔpH-dependent photosynthesis-control in preventing light induced damage to both photosystem II and photosystem I. Sci Rep, 10, 6770.

Bennoun, P., Masson, A. and Delosme, M. (1980) A method for complementation analysis of nuclear and chloroplast mutants of photosynthesis in chlamydomonas. Genetics, 95, 39–47.

Bonente, G., Ballottari, M., Truong, T.B., Morosinotto, T., Ahn, T.K., Fleming, G.R., Niyogi, K.K. and Bassi, R. (2011) Analysis of LhcSR3, a protein essential for feedback de-excitation in the green alga Chlamydomonas reinhardtii. PLoS biology, 9, e1000577.

Burton-Smith, R.N., Watanabe, A., Tokutsu, R., Song, C., Murata, K. and Minagawa, J. (2019) Structural determination of the large photosystem II-light-harvesting complex II supercomplex of Chlamydomonas reinhardtii using nonionic amphipol. The Journal of biological chemistry, 294, 15003–15013.

Chouhan N, Devadasu E, Yadav RM, Subramanyam R. (2022) Autophagy Induced Accumulation of Lipids in pgrl1 and pgr5 of Chlamydomonas reinhardtii Under High Light. Front Plant Sci. Jan 25;12:752634.

Correa-Galvis, V., Redekop, P., Guan, K., Griess, A., Truong, T.B., Wakao, S., Niyogi, K.K. and Jahns, P. (2016) Photosystem II Subunit PsbS Is Involved in the Induction of LHCSR Protein- dependent Energy Dissipation in Chlamydomonas reinhardtii. The Journal of biological chemistry, 291, 17478–17487.

Davis, G.A., Kanazawa, A., Schöttler, M.A., Kohzuma, K., Froehlich, J.E., Rutherford, A.W., Satoh-Cruz, M., Minhas, D., Tietz, S., Dhingra, A. and Kramer, D.M. (2016) Limitations to photosynthesis by proton motive force-induced photosystem II photodamage. eLife, 5.

de Carpentier, F., Lemaire, S.D. and Danon, A. (2019) When Unity Is Strength: The Strategies Used by Chlamydomonas to Survive Environmental Stresses. Cells, 8.

Duysens, L.N. (1954) Reversible Changes in the Absorption Spectrum of Chlorella upon Irradiation. *Science (New York*, N.Y.), 120, 353–354.

Erickson, E., Wakao, S. and Niyogi, K.K. (2015) Light stress and photoprotection in Chlamydomonas reinhardtii. The Plant journal: for cell and molecular biology, 82, 449–465.

Ettinger, W.F., Clear, A.M., Fanning, K.J. and Peck, M.L. (1999) Identification of a Ca2+/H+ antiport in the plant chloroplast thylakoid membrane. Plant physiology, 119, 1379–1386.

Gabilly, S.T., Baker, C.R., Wakao, S., Crisanto, T., Guan, K., Bi, K., Guiet, E., Guadagno, C.R. and Niyogi, K.K. (2019) Regulation of photoprotection gene expression in Chlamydomonas by a putative E3 ubiquitin ligase complex and a homolog of CONSTANS. Proceedings of the National Academy of Sciences of the United States of America, 116, 17556–17562.

Garab, G. and van Amerongen, H. (2009) Linear dichroism and circular dichroism in photosynthesis research. Photosynthesis research, 101, 135–146.

Hendrickson, L., Furbank, R.T. and Chow, W.S. (2004) A simple alternative approach to assessing the fate of absorbed light energy using chlorophyll fluorescence. Photosynthesis research, 82, 73–81.

Im, C.S., Zhang, Z., Shrager, J., Chang, C.W. and Grossman, A.R. (2003) Analysis of light and CO(2) regulation in Chlamydomonas reinhardtii using genome-wide approaches. Photosynthesis research, 75, 111-125.

Järvi, S., Suorsa, M., Paakkarinen, V. and Aro, E.-M. (2011) Optimized native gel systems for separation of thylakoid protein complexes: novel super- and mega-complexes. Biochemical Journal, 439, 207.

Junge, W. and Witt, H.T. (1968) On the ion transport system of photosynthesis--investigations on a molecular level. *Zeitschrift fur Naturforschung. Teil B, Chemie, Biochemie, Biophysik*, Biologie und verwandte Gebiete, 23, 244–254.

Kargul J, Nield J, & Barber J (2003) Three-dimensional reconstruction of a light-harvesting complex I-photosystem I (LHCI-PSI) supercomplex from the green alga Chlamydomonas reinhardtii. Insights into light harvesting for PSI. The Journal of biological chemistry 278(18):16135–16141.

Khona, D.K., Shirolikar, S.M., Gawde, K.K., Hom, E., Deodhar, M.A. and D’Souza, J.S. (2016) Characterization of salt stress-induced palmelloids in the green alga, Chlamydomonas reinhardtii. Algal Research, 16, 434–448.

Kohzuma, K., Cruz, J.A., Akashi, K., Hoshiyasu, S., Munekage, Y.N., Yokota, A. and Kramer, D.M. (2009) The long-term responses of the photosynthetic proton circuit to drought. Plant, cell & environment, 32, 209–219.

Kramer, D.M., Johnson, G., Kiirats, O. and Edwards, G.E. (2004) New Fluorescence Parameters for the Determination of QA Redox State and Excitation Energy Fluxes. Photosynthesis research, 79, 209.

Krause, G.H. (1988) Photoinhibition of photosynthesis. An evaluation of damaging and protective mechanisms. Physiologia Plantarum, 74, 566–574.

Kuroda, H., Kodama, N., Sun, X.Y., Ozawa, S. and Takahashi, Y. (2014) Requirement for Asn298 on D1 protein for oxygen evolution: analyses by exhaustive amino acid substitution in the green alga Chlamydomonas reinhardtii. Plant & cell physiology, 55, 1266–1275.

Laemmli UK (1970) Cleavage of structural proteins during the assembly of the head of bacteriophage T4. Nature 227(5259):680–685.

Laloi, C., Apel, K. and Danon, A. (2004) Reactive oxygen signalling: the latest news. Current opinion in plant biology, 7, 323–328.

Lawlor, D.W. (1970) ABSORPTION OF POLYETHYLENE GLYCOLS BY PLANTS AND THEIR EFFECTS ON PLANT GROWTH. 69, 501–513.

Lemaire, C. and Wollman, F.A. (1989) The chloroplast ATP synthase in Chlamydomonas reinhardtii. II. Biochemical studies on its biogenesis using mutants defective in photophosphorylation. The Journal of biological chemistry, 264, 10235–10242.

Madireddi SK, Nama S, Devadasu E, & Subramanyam R (2019) Thylakoid membrane dynamics and state transitions in Chlamydomonas reinhardtii under elevated temperature. Photosynthesis research 139(1-3):215–226.

Malnoë, A. (2018) Photoinhibition or photoprotection of photosynthesis? Update on the (newly termed) sustained quenching component qH. Environmental and Experimental Botany, 154, 123–133.

Minagawa, J. (2011) State transitions--the molecular remodeling of photosynthetic supercomplexes that controls energy flow in the chloroplast. Biochimica et biophysica acta, 1807, 897-905.

Moroney, J.V., Ma, Y., Frey, W.D., Fusilier, K.A., Pham, T.T., Simms, T.A., DiMario, R.J., Yang, J. and Mukherjee, B. (2011) The carbonic anhydrase isoforms of Chlamydomonas reinhardtii: intracellular location, expression, and physiological roles. Photosynthesis research, 109, 133–149.

Moseley, J.L., Chang, C.W. and Grossman, A.R. (2006) Genome-based approaches to understanding phosphorus deprivation responses and PSR1 control in Chlamydomonas reinhardtii. Eukaryotic cell, 5, 26–44.

Müller, P., Li, X.-P. and Niyogi, K.K. (2001) Non-Photochemical Quenching. A Response to Excess Light Energy. Plant physiology, 125, 1558.

Murata, N., Takahashi, S., Nishiyama, Y. and Allakhverdiev, S.I. (2007) Photoinhibition of photosystem II under environmental stress. Biochimica et biophysica acta, 1767, 414–421.

Nagy, G., Ünnep, R., Zsiros, O., Tokutsu, R., Takizawa, K., Porcar, L., Moyet, L., Petroutsos, D., Garab, G., Finazzi, G. and Minagawa, J. (2014) Chloroplast remodeling during state transitions in Chlamydomonas reinhardtii as revealed by noninvasive techniques in vivo. Proceedings of the National Academy of Sciences of the United States of America, 111, 5042–5047.

Nagy, V., Vidal-Meireles, A., Podmaniczki, A., Szentmihályi, K., Rákhely, G., Zsigmond, L., Kovács, L. and Tóth, S.Z. (2018) The mechanism of photosystem-II inactivation during sulphur deprivation-induced H2 production in Chlamydomonas reinhardtii. The Plant Journal, 94, 548–561.

Naumann, B., Busch, A., Allmer, J., Ostendorf, E., Zeller, M., Kirchhoff, H. and Hippler, M. (2007) Comparative quantitative proteomics to investigate the remodeling of bioenergetic pathways under iron deficiency in Chlamydomonas reinhardtii. Proteomics, 7, 3964–3979.

Nevo, R., Charuvi, D., Tsabari, O. and Reich, Z. (2012) Composition, architecture and dynamics of the photosynthetic apparatus in higher plants. 70, 157–176.

Nilkens, M., Kress, E., Lambrev, P., Miloslavina, Y., Muller, M., Holzwarth, A.R. and Jahns, P. (2010) Identification of a slowly inducible zeaxanthin-dependent component of non-photochemical quenching of chlorophyll fluorescence generated under steady-state conditions in Arabidopsis. Biochimica et biophysica acta, 1797, 466–475.

Nishiyama, Y., Yamamoto, H., Allakhverdiev, S.I., Inaba, M., Yokota, A. and Murata, N. (2001) Oxidative stress inhibits the repair of photodamage to the photosynthetic machinery. EMBO J, 20, 5587–5594.

Niyogi, K.K. (2000) Safety valves for photosynthesis. Current opinion in plant biology, 3, 455–460.

Niyogi, K.K., Shih, C., Soon Chow, W., Pogson, B.J., Dellapenna, D. and Bjorkman, O. (2001) Photoprotection in a zeaxanthin- and lutein-deficient double mutant of Arabidopsis. Photosynthesis research, 67, 139-145.

Peers, G., Truong, T.B., Ostendorf, E., Busch, A., Elrad, D., Grossman, A.R., Hippler, M. and Niyogi, K.K. (2009) An ancient light-harvesting protein is critical for the regulation of algal photosynthesis. Nature, 462, 518.

Petroutsos, D., Busch, A., Janssen, I., Trompelt, K., Bergner, S.V., Weinl, S., Holtkamp, M., Karst, U., Kudla, J. and Hippler, M. (2011) The chloroplast calcium sensor CAS is required for photoacclimation in Chlamydomonas reinhardtii. The Plant cell, 23, 2950–2963.

Petroutsos, D., Tokutsu, R., Maruyama, S., Flori, S., Greiner, A., Magneschi, L., Cusant, L., Kottke, T., Mittag, M., Hegemann, P., Finazzi, G. and Minagawa, J. (2016) A blue-light photoreceptor mediates the feedback regulation of photosynthesis. Nature, 537, 563–566.

Pinnola, A. and Bassi, R. (2018) Molecular mechanisms involved in plant photoprotection. Biochemical Society Transactions, 46, 467.

Porra RJ, Thompson WA, & Kriedemann PE (1989) Determination of accurate extinction coefficients and simultaneous equations for assaying chlorophylls a and b extracted with four different solvents: verification of the concentration of chlorophyll standards by atomic absorption spectroscopy. Biochimica et Biophysica Acta (BBA) - Bioenergetics 975(3):384–394.

Rantala, M., Paakkarinen, V. and Aro, E.M. (2018) Analysis of Thylakoid Membrane Protein Complexes by Blue Native Gel Electrophoresis. Journal of visualized experiments: JoVE.

Rantala, M., Tikkanen, M. and Aro, E.M. (2017) Proteomic characterization of hierarchical megacomplex formation in Arabidopsis thylakoid membrane. The Plant journal: for cell and molecular biology, 92, 951–962.

Rastogi, R. P., Singh, S. P., Häder, D. P., & Sinha, R. P. (2010) Detection of reactive oxygen species (ROS) by the oxidant-sensing probe 2′, 7′-dichlorodihydrofluorescein diacetate in the cyanobacterium Anabaena variabilis PCC 7937. Biochemical and biophysical research communications, 397(3), 603-607.

Schagger H & Vonjagow G (1991) Blue Native Electrophoresis for Isolation of Membrane-Protein Complexes in Enzymatically Active Form. Analytical Biochemistry 199.

Semchonok, D.A., Sathish Yadav, K.N., Xu, P., Drop, B., Croce, R. and Boekema, E.J. (2017) Interaction between the photoprotective protein LHCSR3 and C2S2 Photosystem II supercomplex in Chlamydomonas reinhardtii. Biochimica et Biophysica Acta (BBA) - Bioenergetics, 1858, 379–385.

Sirpiö S, Suorsa M, & Aro EM (2011) Analysis of thylakoid protein complexes by two-dimensional electrophoretic systems. *Methods in molecular biology (Clifton*, N.J.) 775:19–30.

Strenkert, D., Limso, C.A., Fatihi, A., Schmollinger, S., Basset, G.J. and Merchant, S.S. (2016) Genetically Programmed Changes in Photosynthetic Cofactor Metabolism in Copper-deficient Chlamydomonas. The Journal of biological chemistry, 291, 19118–19131.

Subramanyam R, Jolley C, Brune DC, Fromme P, & Webber AN (2006) Characterization of a novel Photosystem I-LHCI supercomplex isolated from Chlamydomonas reinhardtii under anaerobic (State II) conditions. FEBS letters 580(1):233–238.

Takizawa, K., Cruz, J.A., Kanazawa, A. and Kramer, D.M. (2007) The thylakoid proton motive force in vivo. Quantitative, non-invasive probes, energetics, and regulatory consequences of light-induced pmf. Biochimica et biophysica acta, 1767, 1233–1244.

Terauchi, A.M., Peers, G., Kobayashi, M.C., Niyogi, K.K. and Merchant, S.S. (2010) Trophic status of Chlamydomonas reinhardtii influences the impact of iron deficiency on photosynthesis. Photosynthesis research, 105, 39–49.

Tian, L., Nawrocki, W.J., Liu, X., Polukhina, I., van Stokkum, I.H.M. and Croce, R. (2019) pH dependence, kinetics and light-harvesting regulation of nonphotochemical quenching in Chlamydomonas. Proceedings of the National Academy of Sciences of the United States of America, 116, 8320–8325.

Tokutsu, R., Fujimura-Kamada, K., Matsuo, T., Yamasaki, T. and Minagawa, J. (2019) The CONSTANS flowering complex controls the protective response of photosynthesis in the green alga Chlamydomonas. Nat Commun, 10, 4099–4099.

Tokutsu, R. and Minagawa, J. (2013) Energy-dissipative supercomplex of photosystem II associated with LHCSR3 in <em>Chlamydomonas reinhardtii</em>. Proceedings of the National Academy of Sciences, 110, 10016.

Velthuys, B.R. (1978) A third site of porton translocation in green plant photosynthetic electron transport. Proceedings of the National Academy of Sciences of the United States of America, 75, 6031–6034.

Wittig, I., Braun, H.-P. and Schägger, H. (2006) Blue native PAGE. Nature Protocols, 1, 418–428.

Yruela, I. (2013) Transition metals in plant photosynthesis. Metallomics, 5, 1090-1109.

Zhang, Z., Shrager, J., Jain, M., Chang, C.W., Vallon, O. and Grossman, A.R. (2004) Insights into the survival of Chlamydomonas reinhardtii during sulfur starvation based on microarray analysis of gene expression. Eukaryotic cell, 3, 1331–1348.

