## supplementary for "Distinctive mechanism of LHCSR3 expression and function under osmotic stress in *Chlamydomonas reinhardtii*"

**This PDF file includes:**

Supplementary text
Figures S1 to S3

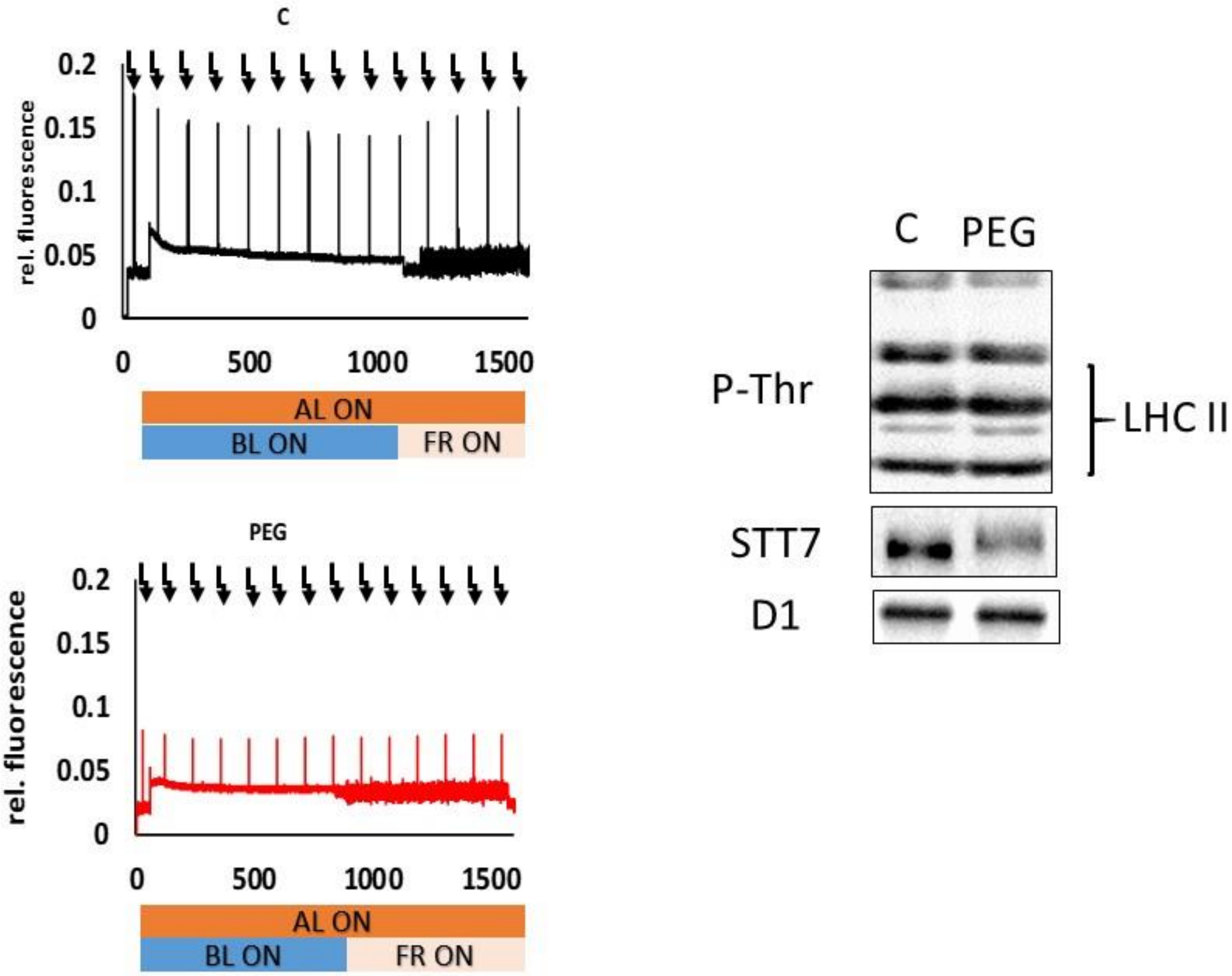


**Figure S1. Analysis of state transitions through Chlorophyll *a* fluorescence kinetics in *C. reinhardtii* cells grown under osmotic stress.** A) *C. reinhardtii* (Control (C) and osmotically stressed (PEG)) cells were brought to State I by vigorous agitation (~200rpm) in the dark for 30 min, and Fv/Fm values were determined. State II was induced by illumination with actinic light (AL) supplemented with blue light (BL). Afterwards, State I was induced by supplementing actinic light with far-red light (FL). During actinic light treatment saturating pulses were applied every 2 min to follow Fm` kinetics. B) Immunoblot analysis of LHCII phosphorylation, Stt7 content from the cells grown under control (C) and osmotic stress (PEG).

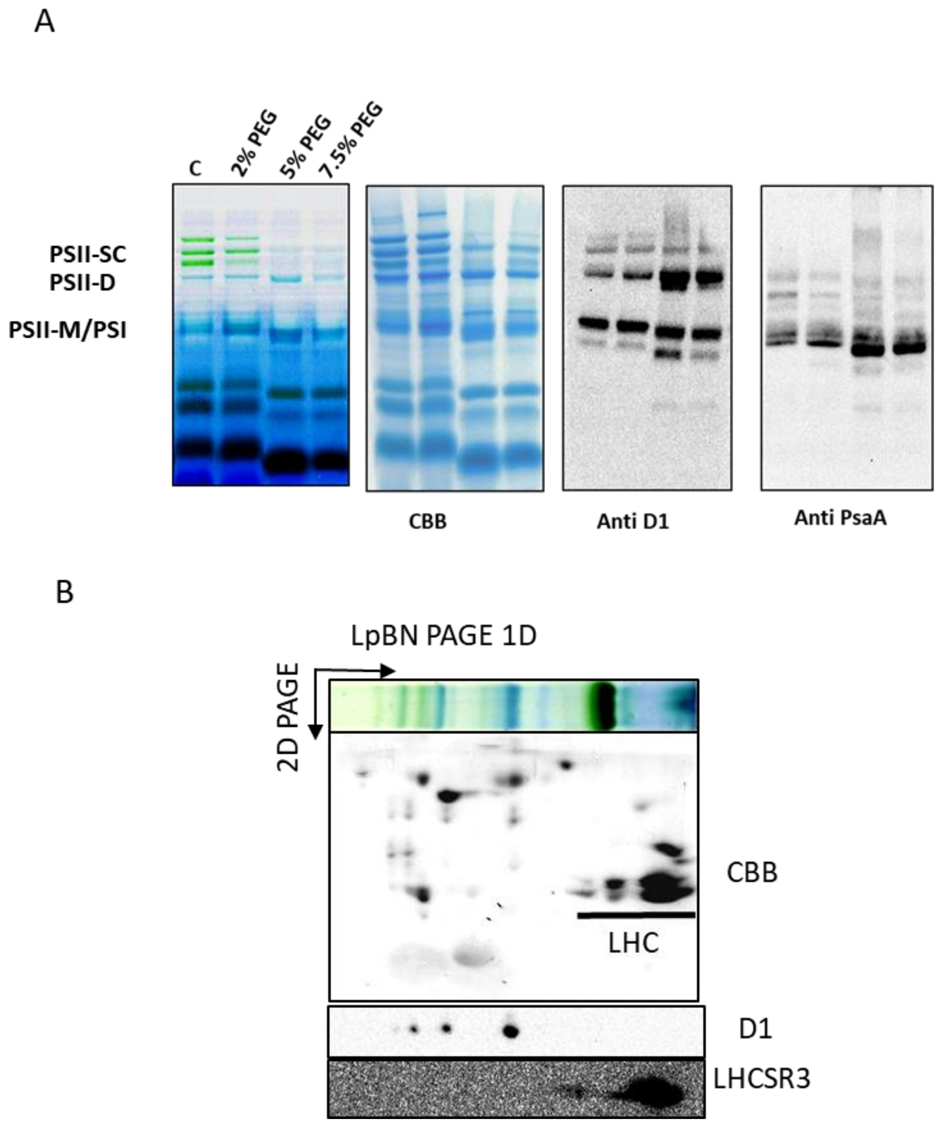


**Figure S2**. **Identification of Blue native gel bands from *C. reinhardtii* cells grown in different concentrations of PEG.** A) PSII-SC photosystem II supercomplex, PSII-D photosystem II dimer, PSII-M photosystem II monomer, PSI photosystem I. As indicated, the BN-PAGE gel was used for immunoblotting against D1, PsaA and PetB antibodies. B) Detection of LHCSR3 localization by immunoblotting of 2D/BN SDS gel of thylakoids from PEG-grown cells; immunoblotting is performed using anti-D1 and anti-LHCSR3 antibodies.

A

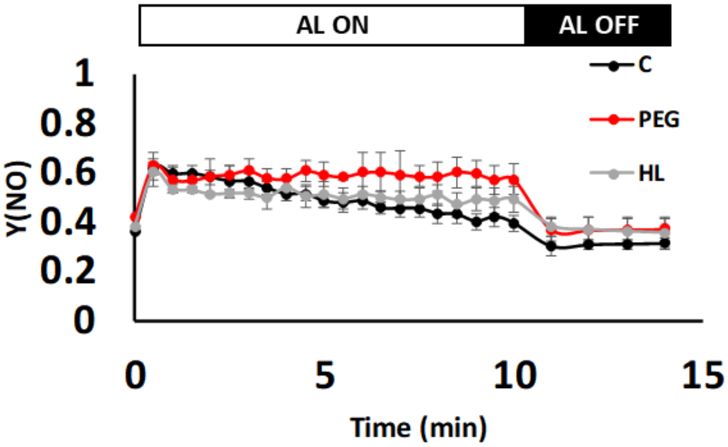
AL ON
AL OFF
C
PEG
HL
Y(NO)
1
0.8
0.6
0.4
0.2
0
0
5
10
15
Time (min)

B

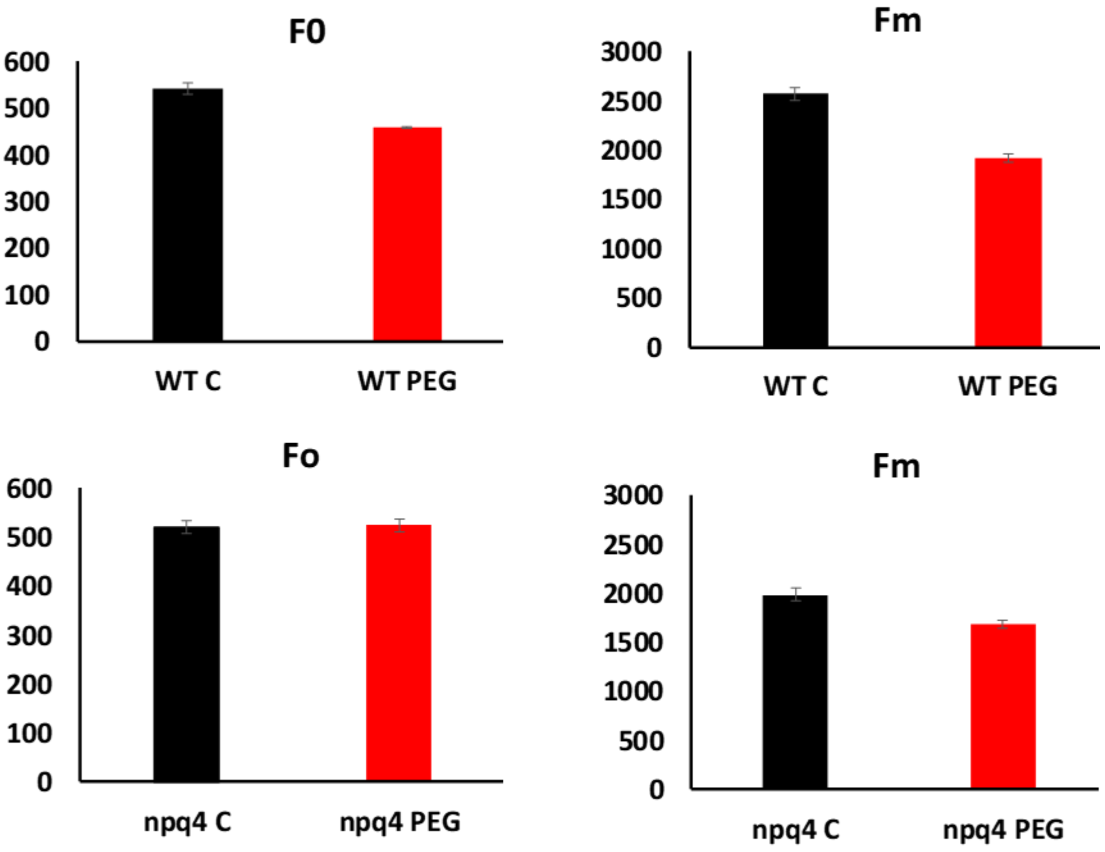
F0
600
500
400
300
200
100
0
WT C
WT PEG
Fm
3000
2500
2000
1500
1000
500
0
WT C
WT PEG
Fo
600
500
400
300
200
100
0
npq4 C
npq4 PEG
Fm
3000
2500
2000
1500
1000
500
0
npq4 C
npq4 PEG

**Figure S3. Kinetics of Non-regulated quenching and evidence of sustained quenching under osmotic stress.** A) Induction of non-regulated energy loss Y(NO) of the cells grown under different conditions. Values represent means ± SD (n = 3). B) Comparison of changes in minimal and maximal fluorescence of WT and npq4 mutant under osmotic stress. Values represent means ± SD (n = 5).
